# Structural and Dynamic Insights into the Biased Signaling Mechanism of the Human Kappa Opioid Receptor

**DOI:** 10.1101/2024.04.11.588819

**Authors:** Chiyo Suno-Ikeda, Ryo Nishikawa, Riko Suzuki, Seiya Iwata, Tomoyo Takai, Takaya Ogura, Mika Hirose, Akitoshi Inoue, Eri Asai, Ryoji Kise, Yukihiko Sugita, Takayuki Kato, Hiroshi Nagase, Tsuyoshi Saitoh, Kota Katayama, Asuka Inoue, Hideki Kandori, Takuya Kobayashi, Ryoji Suno

## Abstract

The κ-opioid receptor (KOR) is a member of the G protein-coupled receptor (GPCR) family, responsible for modulating cellular responses through transducers such as G proteins and arrestins. G protein-biased KOR agonists hold promise due to their potential to mitigate side effects such as drug aversion and sedation while preserving analgesic and antipruritic effects. Here, we shed light on the structural dynamics of the human KOR-G_i_ signaling complex bound with either nalfurafine (a G-protein-biased agonist) or U-50,488H (a balanced agonist) using cryo-electron microscopy (cryo-EM). Cryo-EM structures of the KOR-G_i_ signaling complexes identify the ligand binding mode in the activated state. Vibrational spectroscopy analysis reveals changes in the ligand-binding pocket upon binding to these ligands. Cell-based mutant experiments pinpoint four amino acids (K227^5.40^, C286^6.47^, H291^6.52^, and Y312^7.34^; Ballesteros–Weinstein numbering is shown in superscript) that play crucial roles in arrestin recruitment. Among these four amino acids, H291^6.52^ and Y312^7.34^ are also implicated in G-protein coupling. Our findings pave the way for targeting specific residues in the KOR ligand-binding pocket to enhance KOR-mediated therapeutic effects while mitigating unwanted side effects.

## INTRODUCTION

G protein-coupled receptors (GPCRs) mediate intracellular processes by interacting with signaling transducers such as G proteins and arrestins, a process initiated by binding with extracellular agonists^1^. Opioid receptors, members of the GPCR superfamily, are categorized into four subclasses: μ, δ, κ, and nociceptin receptors (MOR, DOR, KOR, and NOP receptors, respectively). These receptors are recognized for their role in mediating the analgesic properties of opioid molecules^2^.

Upon agonist binding, opioid receptors activate the G_i/o_ subtype of G protein family, initiating a cascade of intracellular signaling pathways. This process attenuates the excitation of pain-sensing neurons and induces analgesia^3^. Studies with striatal neurons from arrestin knockout mice have revealed the involvement of the arrestin pathway in mediating adverse effects, including drug aversion and sedation^4–6^. Consequently, there has been a concerted effort to develop G protein-biased agonists with minimal arrestin-recruitment activity, aiming to create analgesics devoid of adverse side effects^7,8^. MOR agonists like morphine and fentanyl exhibit high analgesic efficacy but are linked with side effects such as respiratory depression and drug dependence. G-protein biased agonists such as TRV130, PZM21, and SR-17018 were developed to mitigate these side effects^2^. Despite these innovations, fully suppressing adverse effects remains elusive. Agonists for KOR and DOR also lead to side effects like sedation and convulsions/catalepsy, respectively^9^. Several G protein-biased ligands for DOR, including derivatives of SNC80 (e.g., ARM390), KNT-127 and those based on the morphinan backbone (SB-235863), have been developed to mitigate side effects^9,10^. Particularly, KNT-127 reduces the side effects of convulsions but is not approved for analgesia^11^. Similarly, for KOR, compounds like 6’-GNTI, nalfurafine^12^, novocaine, triazole 1.1 and its derivatives, salvinorin A derivatives, and GR89696 have been developed as G protein biased agonists. While nalfurafine, approved as an antipruritic drug, successfully eliminates the side effect of drug aversion, side effect of sedation persists at doses showing analgesic effects.

Despite the development of biased agonists, the intricacies of how these compounds cause selective signaling of GPCRs remain largely unexplored^2^. To comprehend the mechanism of action of drugs, the structures of opioid receptors bound to drugs with various properties have been determined. Further detailed analyses, including molecular dynamics simulations and pharmacological studies, are revealing biases in ligand binding modes and mechanisms of action^13^.

With respect to the structural biology of KOR, the X-ray crystal structure of KOR bound to nalfurafine and a G-protein mimicking nanobody (Nb39) was recently determined^14^. Comparison with molecular dynamics simulations of KOR in the nalfurafine, U-50,488H and arrestin signalling selective agonist WMS-X600 bound states revealed the presence of three active states. Molecular dynamics simulations and pharmacological analysis of KORs bound with G-protein/arrestin-biased and balanced agonists have demonstrated that the orientation of the side chain of Q115^2.60^ and the distance between K227^5.40^ and E297^6.58^ differ for each ligand-bound state, thereby affecting signal selectivity. However, no comparison between the structures of the G-protein-biased and balanced agonist binding states has been conducted, necessitating further studies. Additionally, experimental observations of agonist-dependent dynamic conformational changes in KORs prior to the binding of signaling molecules have not been made, and the information on selective signaling mechanisms is still developing. In this study, we present cryo-EM structures of the KOR-G_i_ signaling complexes bound with the balanced agonist U-50,488H and the G protein-biased agonist nalfurafine. Furthermore, we analyze ligand-dependent dynamic changes in the amino acid side chains of KOR using attenuated total reflection-Fourier transform infrared spectroscopy (ATR-FTIR), a type of vibrational spectroscopy. Based on the structural information and the spectroscopy dynamics data, we conducted a mutagenesis study and identified amino acid residues (F227^5.40^, C286^6.47^, H291^6.52^, Y312^7.34^) that play crucial roles in each signal transduction pathway.

## RESULTS

### Overall structures of nalfurafine- or U-50,488H-bound KOR-G_i_ complexes

The cryo-EM structures of nalfurafine- or U-50,488H-bound human KOR-G_i_ signaling complexes were determined at 2.76 Å and 2.9 Å resolution from 858,423 and 1,225,096 particles, respectively (Supplemental Figs. 1a-d, 2a-2c and 3a-3c). This allowed us to precisely identify and assign the transmembrane (TM) domain of KOR, the G_i_ protein heterotrimer, the antibody fragment, and ligands in the cryo-EM map (Figs. 1a, 1b, 1c, and Supplementary Figs. 4a and 4b). Overlaying the receptor regions of the nalfurafine- and U-50,488H-bound KOR-G_i_ complexes revealed a close alignment with a backbone root mean square deviation (RMSD) of 0.356 Å. Analysis of the cryo-EM data from the nalfurafine-bound KOR-G_i_ complex yielded four different maps, of which the highest resolution map was used for structure building and final refinement. Modeling all four maps and examining their differences revealed a subtle rotation of the G protein relative to the receptor, suggesting that G_i_ can form several dynamic binding states to the KOR (Supplementary Fig. 2d).

**Figure 1:**
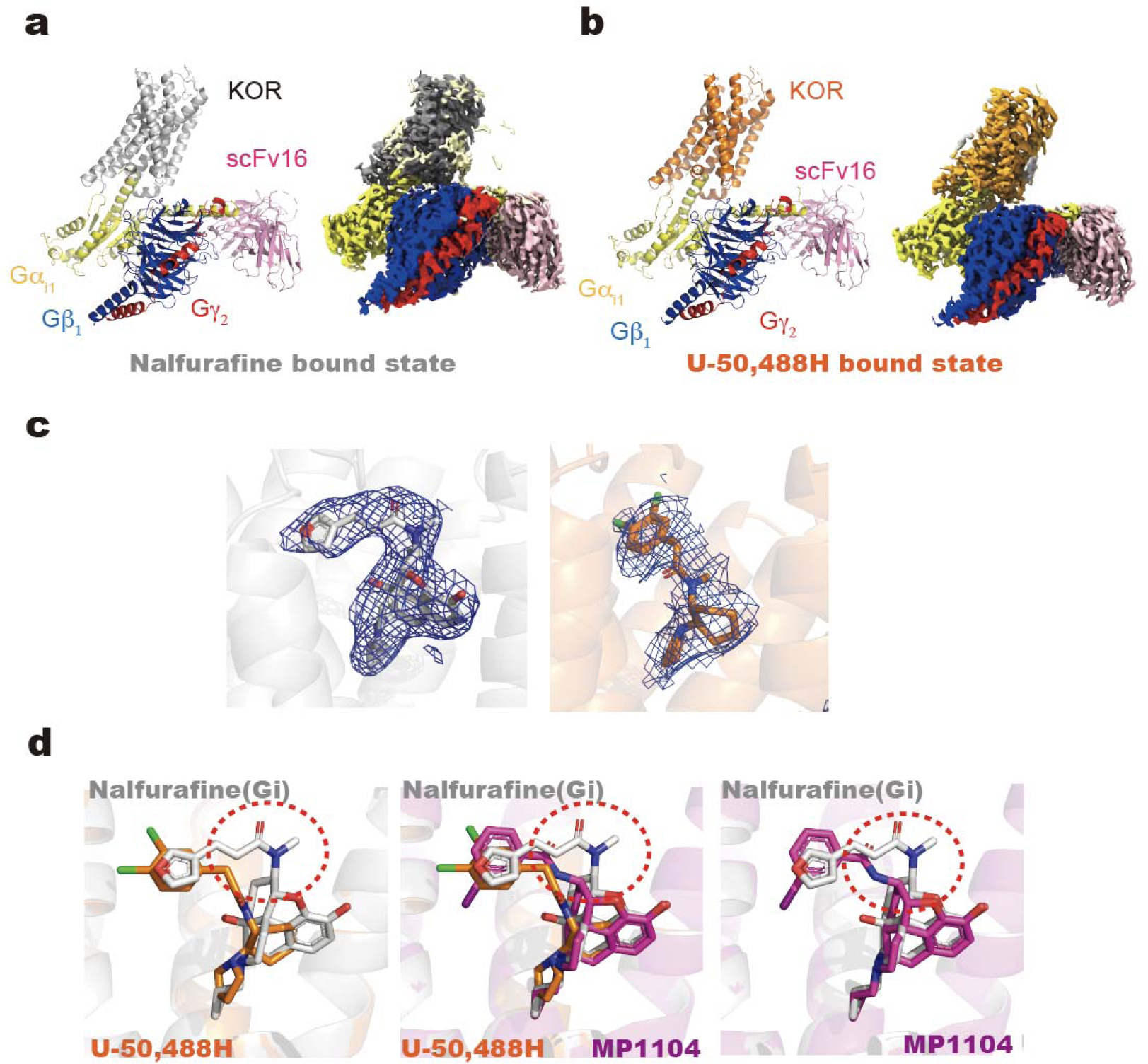
Cryo-EM structures of the biased/balanced agonist-bound KOR-Gi signaling complexes. Comparison of the structural model with the cryo-EM density of the entire complex. Structures and cryo-EM maps of the KOR-Gi complex in nalfurafine- (a) and U-50,488H-bound (b) states. Nalfurafine-bound KOR is gray, U-50,488H-bound KOR is displayed orange, Gi is yellow, Gβ is blue, Gγ is red, and scFv16 is pink. (c) Cryo-EM density maps and models of of nalfurafine (left) or U-50,488H (right). Maps are shown in blue. (d) Differences in ligand binding mode of nalfurafine (gray), U-50,488H (orange), and MP1104(magenta). The red dotted circles indicate characteristic regions of nalfurafine’s structure compared to other agonists.

To elucidate the differences in the binding modes of G protein-biased and balanced agonists, we compared the structures of the two KOR-G_i_ complexes determined in this study with the previously reported MP1104 (a non-selective balanced agonist of opioid receptors)-bound state^15^. The major difference observed in ligand-binding mode was the interaction of nalfurafine with TM5, particularly K227^5.40^. In contrast, U-50,488H and MP1104 exhibited no interaction with K227^5.40^ (Figs. 1c and 2a).

**Figure 2:**
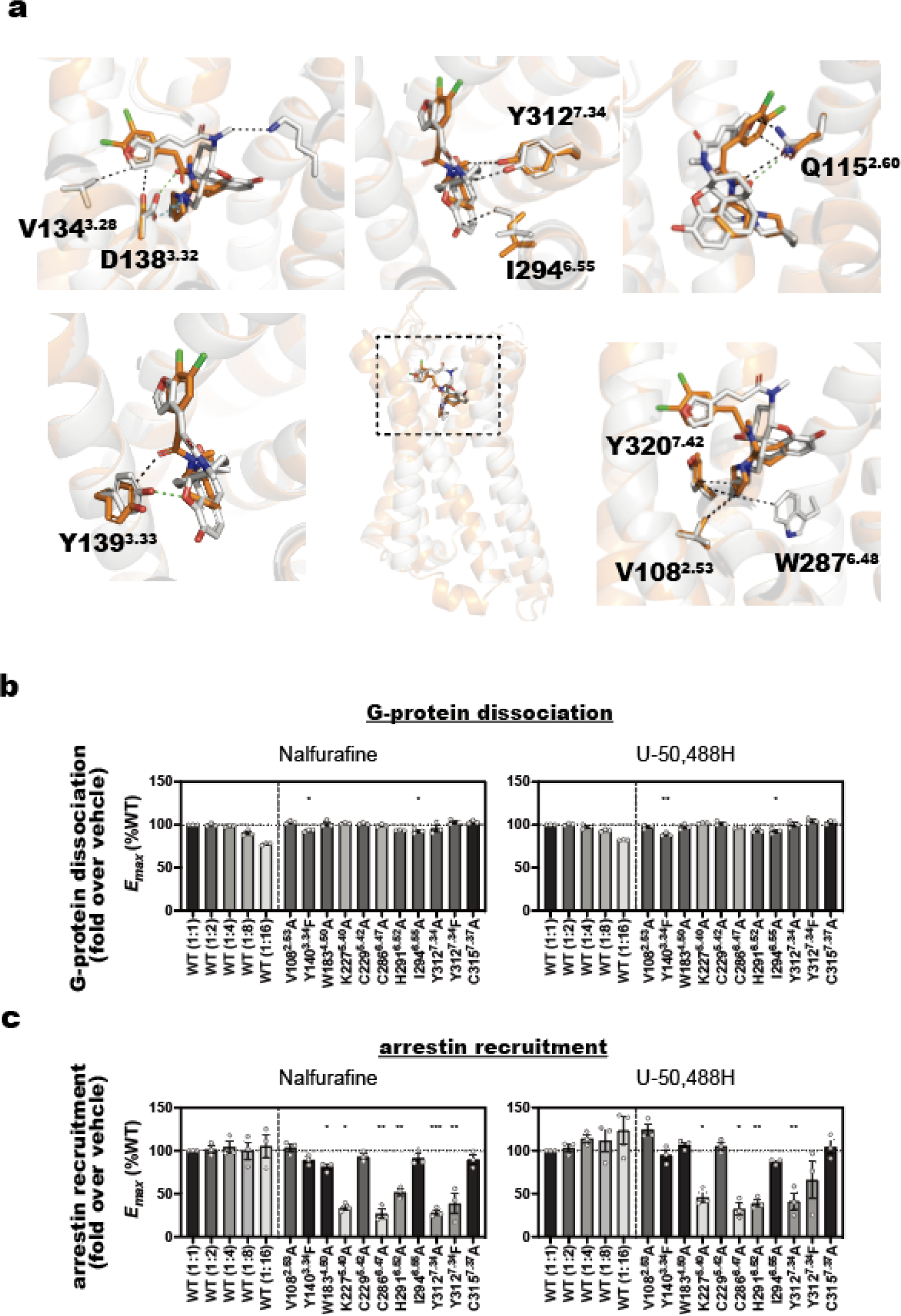
Different binding modes of KOR-biased/balanced agonists. Amino acid residues of KOR that interact with nalfurafine or U-50,488H (a). Van der Waals interactions are represented by black, green, and cyan dotted lines for hydrogen and ionic bonds, respectively. Amino acids that are within 4 Å of the ligand are displayed as sticks. Nalfurafine-bound KOR is gray, U-50,488H-bound KOR is orange. Gi-coupling activity analyzed by the NanoBiT-G-protein dissociation assay (b) and β-arrestin2-recruiting activity analyzed by the NanoBiT-β-arrestin recruitment assay (c). Data are presented as mean values ± SEM (n = 3-4; dots). For individual experiments performed in parallel, data were normalized to the wild-type KOR (1:1) and presented as Emax. The colors in the mutant bars correspond to the expression-matched wild-type conditions (Supplementary Figure 5).

### Comparison of the Ligand-Binding Modes of KOR: Biased vs. Balanced Agonists

Structural determination of the KOR-G_i_ signaling complexes in nalfurafine- and U-50,488H-bound forms unveiled the details of their respective agonist binding modes (Fig. 2a). Key residues, including K227^5.40^, W287^6.48^, and I294^6.55^ located in TM 5 and 6, exhibited exclusive van der Waals interactions with nalfurafine, while no interaction with U-50,488H was observed. In TM2, both agonists formed van der Waals interactions with the side chains of V108^2.53^; the side chain of Q115^2.60^ formed a van der Waals interactions with U-50,488H and a hydrogen bond with nalfurafine. In TM3, the side chain of V134^3.28^ formed a van der Waals interaction only with nalfurafine. The side chain of D138^3.32^ forms van der Waals interactions with U-50,488H and forms hydrogen bond and ionic bond with nalfurafine. Similarly, the side chain of Y139^3.33^ interacts with U-50,488H via van der Waals forces but forms hydrogen bonds with nalfurafine. In TM7, both Y312^7.34^ and Y320^7.42^ interact with the agonists through van der Waals interactions. The higher affinity of nalfurafine compared to U-50,488H may be attributed to the greater number of interacting amino acids with nalfurafine. Furthermore, the interaction with the side chains of D138^3.32^ and Y139^3.33^ is stronger for nalfurafine due to hydrogen and ionic bonding, contrasting with U-50,488H, which relies on van der Waals interactions.

To validate the functional importance of agonist-interacting residues, we conducted a mutagenesis study using NanoBiT-based G protein-dissociation assay and arrestin-recruitment assay. Mutants were chosen based on amino acids located within less than 4 Å from the ligand, as depicted in Fig. 2a. The expression levels of mutants were confirmed by flow cytometry (Supplemental Fig. 5). Overall, neither of the ligands exhibited a significant effect on the *E_max_* in G protein activity, although there were slight changes in EC_50_ (Fig. 2b, Supplementary Fig. 6, and Supplementary Table 1). However, K227^5.40^A and Y312^7.34^A/F showed significantly reduced arrestin activity for both ligands (Fig. 2c and Supplementary Fig. 8).

### Identification of crucial amino acid residues for arrestin recruitment

Two pivotal amino acids influencing arrestin recruitment were examined through a structural comparison between nalfurafine- and U-50,488H-bound KOR-G_i_ signaling complexes, alongside signal assays using their respective agonists. First, we focused on K227^5.40^, which exclusively binds to nalfurafine and affects signaling activity of U-50,488H. Comparative structural analysis of the two KOR-G_i_ signaling complexes revealed direct interaction of K227^5.40^ side chain with nalfurafine but not with U-50,488H. In the U-50,488H-bound KOR structure, a weak ionic bond was observed between the side chains of K227^5.40^ and E297^6.58^, absent in the nalfurafine-bound KOR structure (Fig. 3a).

**Figure 3:**
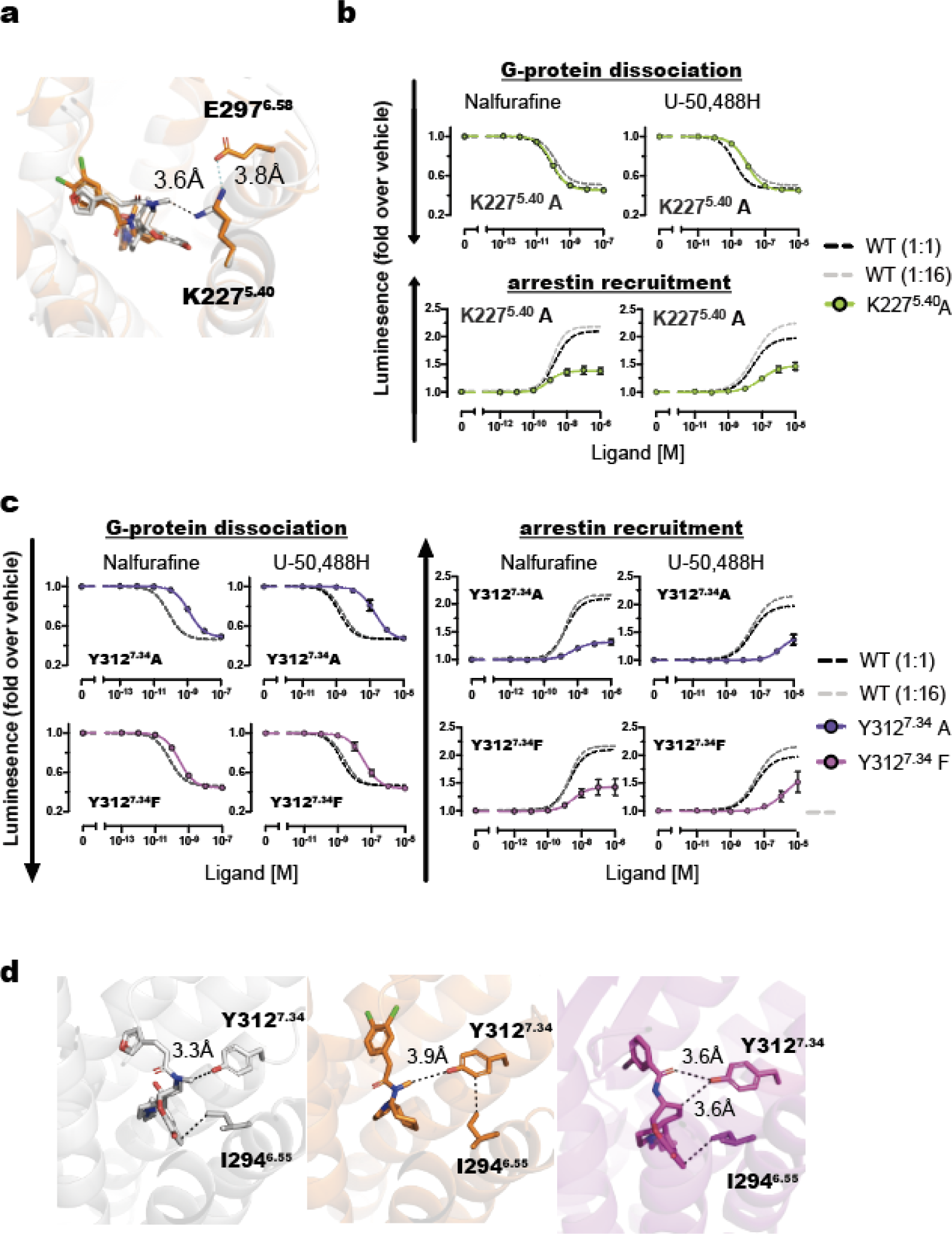
Contribution of K2275.40 and Y3127.34 to the signal selectivity of KOR. Binding mode of the side chain of K227^5.40^ to the agonists (a). Van der Waals interactions are depicted in black dotted lines, while hydrogen bonds and ionic bonds are represented by green dotted lines. Amino acids that are within 4 Å of the ligand are displayed as sticks. Concentration-response curves of the NanoBiT-G-protein dissociation assay and the NanoBiT-β-arrestin recruitment assay of Y227^5.40^ (b) and Y312^7.34^ (c) mutants. Dashed lines in the mutant panels represent the wild-type KOR (1:1) response. Data are presented as mean values ± SEM (n = 3). Note that in numerous data points, the error bars are smaller than the size of the symbols, making them not visible. Binding mode of the side chain of Y312^7.34^ to the agonists. Binding mode of the side chain of Y312^7.34^ to the agonists (d). Nalfurafine-bound KORs are represented in gray, U-50,488H-bound KORs is orange, and MP-1104-bound KOR is magenta.

In the NanoBiT assays, the K227^5.40^A mutation did not affect the *E_max_* of G protein coupling but significantly reduced the *E_max_* of arrestin-recruitment activity in the presence of both agonists. Despite the lack of direct interaction between K227^5.40^ and U-50,488H, the K227^5.40^A mutation notably reduced arrestin recruitment, emphasizing the pivotal role of the K227^5.40^-E297^6.58^ interaction in this process. Conversely, K227^5.40^A exhibited largely unchanged G protein coupling with nalfurafine, albeit with reduced pEC_50_ in the presence of U-50,488H (Fig. 3b, Supplementary Figs. 6-9, and Supplementary Table 1).

Essentially, interaction with K227^5.40^ proved dispensable for the G-protein response to nalfurafine, highlighting its exclusive relevance to arrestin recruitment. These findings confirm nalfurafine’s biased signaling activity, inhibiting the K227^5.40^-E297^6.58^ interaction by interacting with the side chain of K227^5.40^ and consequently diminishing arrestin-recruitment activity while maintaining G-protein coupling. Molecular dynamics simulations proposed by Daibani *et al.* ^14^ support the significance of the K227^5.40^-E297^6.58^ interaction in arrestin recruitment, aligning with our structural insights and pharmacological analyses.

Next, we investigated the Y312^7.34^A and Y312^7.34^F mutants, which showed a significant reduction in the *E_max_* of arrestin-recruitment activity (Fig. 3c, Supplementary Figs. 6-9, and Supplementary Table 1). Specifically, the Y312^7.34^A mutant exhibited a markedly decreased pEC_50_ for G protein-coupling activity and a substantially lower *E_max_* for arrestin-recruitment activity with both ligands. Although the G-protein-coupling activity of the Y312^7.34^F mutant with nalfurafine resembled that of the wild type, its pEC_50_ with U-50,488H decreased. These findings imply that the hydroxyl group of the Y312^7.34^ side chain plays a crucial role in nalfurafine-mediated arrestin-recruitment activity.

The distances from the hydroxyl group on the side chain of Y312^7.34^ to U-50,488H and nalfurafine measured at 3.9 Å and 3.3 Å, respectively. The absence of a hydroxyl group in the Y312^7.34^F mutant increases the distance between the phenylalanine side chain and the ligand, weakening the affinity. KOR-G_i_ signaling structures shows that the interaction between these two agonists and the side chain of Y312^7.34^ involves van der Waals interactions, and similar van der Waals interactions are expected with the Y312^7.34^F phenylalanine side chain. This finding suggests that the appropriate distance between the side chain of Y312^7.34^ and the agonist, determining the strength of interaction, influences the signaling activity of G proteins and arrestins.

For nalfurafine, the distance at which it interacts with the side chain of Y312^7.34^ was sufficient to demonstrate G protein coupling activity but relatively unfavorable for arrestin-recruitment activity. In the KOR structure bound to MP1104, the intermolecular distance between the side chain of Y312^7.34^ and MP1104 is 3.6 Å, which falls between the distances from this side chain to nalfurafine and U-50,488H. Given that MP1104 is a balanced agonist, this suggests that the distance between the ligand and Y312^7.34^ is stringent for properties regulating arrestin recruitment (Fig. 3d).

Based on molecular dynamics simulations, Daibani *et al.* identified Q115^2.60^ as an amino acid involved in KOR signaling^14^. To understand the role of Q115^2.60^, we compared the structures of KOR-G_i_ signaling complexes bound to nalfurafine or U-50,488H (Supplementary Fig. 10). In the KOR structure bound to U-50,488H, the side chain of Q115^2.60^ engaged in a weak van der Waals interaction with the ligand and formed a hydrogen bond with the side chain of Y320^7.42^. Conversely, nalfurafine interacted with the side chain of Q115^2.60^ at two locations, establishing both a van der Waals interaction and a hydrogen bond. This suggests that nalfurafine forms a stronger interaction with the side chain of Q115^2.60^ than U-50,488H. Previous MD simulations have indicated that when KOR binds to WMS-X600, an arrestin-biased agonist, the side chain of Q115^2.60^ orient towards both Y320^7.42^ and Y66^1.39^. These observations suggest that differences in agonist binding with Q115^2.60^ result in different side chain orientations of Q115^2.60^, which affect arrestin-recruitment activity (Supplementary Fig. 10).

### Quantifying agonist-dependent conformational changes in KOR using infrared spectroscopy

The active state of KOR bound to G-protein, as determined by cryo-EM, unveils the respective agonist binding mode, albeit post-G-protein binding. The signal selectivity of biased agonists for KOR may aid in structuring environments conducive to the binding of each signaling transducer. Next, we examined the conformational changes observed upon binding of G-protein-biased/balanced agonists to KORs before the binding of signal transducers, utilizing ATR-FTIR spectroscopy^16–18^ (see Figs. 4a and 4b).

**Figure 4.**
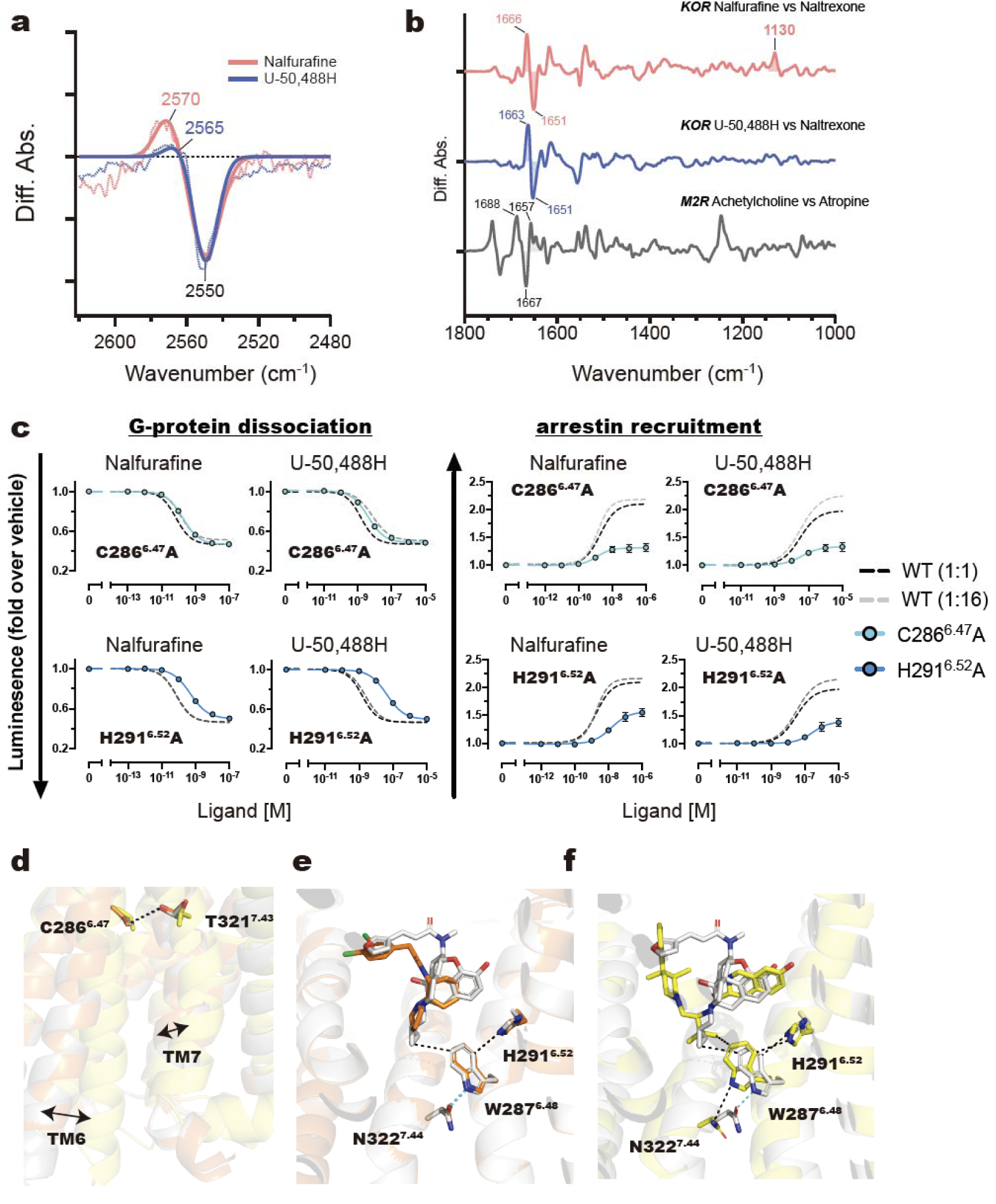
Ligand binding-dependent conformational changes detected by ATR-FTIR and their effect on signal selectivity. FTIR spectroscopy was employed to detect conformational changes in the KOR upon agonist binding. Signals from nalfurafine and U-50,488H were observed as they interacted with the KOR in the presence of the antagonist naltrexone. ATR FTIR spectra illustrating the conformational changes in cysteine (a) and histidine (b). (c) Concentration-response curves of the NanoBiT-G protein dissociation assay and the NanoBiT-β-arrestin recruitment assay of C286^6.47^A and H291^6.52^A mutants. Dashed lines in the mutant panels represent the wild-type (WT) KOR (1:1) response. Data are presented as mean values ± SEM (n = 3). Note that in numerous data points, the error bars are smaller than the size of the symbols, making them not visible. (d) Interaction between C286^6.47^ and T321^7.43^ and conformational changes in TM6 during agonists and antagonist binding. Arrows indicate the extent of conformational change in TM6 when comparing inactive and active forms. (e) Interaction network formed between agonists, H291^6.52^ and N322^7.44^, which links with a toggle switch (W287^6.48^) that is common to GPCRs and important for their activation. (f) Interaction of H291^6.52^ and W287^6.48^ in agonist- or antagonist-bound states. Nalfurafine-bound KORs are represented in gray, U-50,488H-bound KORs is in orange, and inverse agonist JDTic-bound KORs are displayed in magenta.

Upon introducing both nalfurafine and U-50,488H to the inactive state of KOR bound with the antagonist naltrexone, distinct changes in the infrared signals emerged at amide-I of an α-helix (1,700–1,600 cm^-1^), C-N stretching of histidine imidazole (1,200–1,100 cm^-1^), and S-H stretching of cysteine side chain (2,600–2,500 cm^-1^) vibration regions. The band around 2,550 cm^-1^, characteristic of the S-H stretching vibration of cysteine, exhibited a band-shift for both ligand-bound forms, indicating a difference in the hydrogen-bonding environment of cysteine (refer to Fig. 4a). In Fig. 4b, the amide-I band up-shifted from 1,651 cm^-1^ to 1,666 cm^-1^ and 1,663 cm^-1^ upon ligand exchanging from naltrexone to nalfurafine and U-50,488H, respectively. This result suggests that the α-helix underwent perturbation during the conformational transition from the antagonist-bound inactive state to the agonist-bound active state.

Previously, a similar ligand binding-induced conformational change analysis was reported for the muscarinic receptor (M_2_R)^17,18^. A comparison with KOR, however, revealed that the amide-I shift between KOR and M_2_R is opposite (see Fig. 4b) (KOR:1,666 or 1663 cm-1 (agonist bound form) --> 1,651 cm-1 (antagonist bound form) spectral up-shift, M_2_R: 1,667 cm^-1^ (antagonist bound form) --> 1,657 cm^-1^ (agonist bound form) spectral down-shift). While it is generally understood that class A GPCRs undergo significant movement of TM6 upon agonist binding, there may be diversity in local α-helical perturbation at TM6 among GPCRs.

Conversely, in Fig. 4b, the negative bands of the naltrexone-bound form are identical to the red and blue lines, yet a 3 cm^-1^ variance is observed between the nalfurafine-bound form (red line: 1,666 cm^-1^) and the U-50,488H-bound form (blue line: 1,663 cm^-1^). This disparity suggests that G-protein-biased ligand (nalfurafine) binding induces more local α-helical perturbation. Furthermore, the positive 1,130 cm^-1^ band likely reflects the characteristic C-N stretching vibration of histidine imidazole^19^ and is amplified upon binding of nalfurafine.

Cysteine residues in the KOR involved in signaling activity were examined through mutant experiments. Structural information of KOR reveals three cysteines (C229^5.41^, C286^6.47^, and C315^7.37^) that do not directly interact with the agonist. These cysteine residues were substituted with alanine, and their impact on signaling activity was assessed. The C229^5.41^A mutation exhibited no discernible change in G protein binding and arrestin-recruitment. However, the C286^6.47^A mutant displayed a considerable decrease in *E_max_* of arrestin-recruitment activity for both ligands, while the C315^7.37^A mutant exhibited a decrease in G protein pEC_50_ and a slight decrease in arrestin-recruitment activity only in the presence of U-50,488H (Figs. 4c, Supplementary Figs. 6–9, and Supplementary Table 1). Specifically focusing on C286^6.47^A, which demonstrated a significant change in signaling activity, no observable interactions were noted between the side chains of C286^6.47^ and T321^7.43^ in the inactive state, suggesting that these interactions are promoted in an agonist binding-dependent manner (Fig. 4d).

The mutation C286^6.47^A resulted in a greater distance from the side chain of T321^7.43^ compared to the wild type, indicating a weakened interaction. This observation, coupled with the mutagenesis analysis, suggests that the interaction between the side chains of C286^6.47^ and T321^7.43^ is essential for arrestin-recruitment activity. Specifically, the G protein-coupling activity of C286^6.47^A was nearly identical to that of the wild type upon the addition of nalfurafine, implying that this interaction does not impact G-protein dissociation. Regarding C315^7.37^, its side chain interacts with the oxygen of the main chain carbonyl group of P289^6.50^ (Supplementary Fig. 11). In the presence of U-50,488H, both the G-protein dissociation and arrestin recruitment activities of C315^7.37^A were slightly impaired (Supplementary Figs. 6–9 and Supplementary Table 1). These findings suggest that the interaction of C315^7.37^ side chain with the oxygen of the P289^6.50^ carbonyl group is crucial for forming a pocket suitable for U-50,488H binding. Next, histidine residues in KOR associated with signaling activity were investigated through mutagenesis. Two histidine residues of KOR, H162^ICL^^2^ and H291^6.52^, were selected for structural analysis. KOR-G_i_ signaling structures shows that the side chain of H162^ICL^^2^ remained exposed in solution without interaction with any amino acid side chain or ligand. Conversely, structural data from the KOR-G_i_ complex in the nalfurafine-bound state confirmed an interaction between the side chains of H291^6.52^ and W287^6.48^ (Figs. 4e and 4f).

The mutagenesis assay revealed that the H291^6.52^A mutant reduced both G-protein dissociation and arrestin recruitment activities. Notably, the decrease in *E_max_* of arrestin-recruitment activity highlights the importance of side-chain interactions between W287^6.48^ and H291^6.52^ in the arrestin response (Fig. 4c, Supplementary Figs. 6-9, and Supplementary Table 1). A noticeable rotation of the side chain of H291^6.52^ was observed in the agonist-bound state of KOR in comparison to the inverse agonist JDTic-bound state (Fig. 4f), which was identified as a signal in the infrared spectroscopy.

In the nalfurafine-bound state, W287^6.48^ directly interacts with the ligand, while H291^6.52^ shows no direct interaction with either ligand. This suggests that H291^6.52^ is crucial for stabilizing the orientation of the W287^6.48^ side chain, indirectly influencing ligand binding and signaling activity (Figs. 4e and 4f). Despite the distance between U-50,488H and W287^6.48^ being more than 4 Å, both W287^6.48^A and H291^6.52^A mutants significantly reduced arrestin-recruitment activity like the addition of nalfurafine, indicating a molecular mechanism for arrestin recruitment similar to nalfurafine.

Furthermore, H291^6.52^ in TM6 interacts with W287^6.48^, forming an interaction network with N322^7.44^ of TM7. These interactions underscore the significant role played by the interplay between TM6 and TM7 in the arrestin-recruitment activity of KOR (Figs. 4e, 4f, Supplementary Figs. 6-9, and Supplementary Table 1).

ATR FTIR spectra illustrating the conformational changes in cysteine (a) and histidine (b). (c) Concentration-response curves of the NanoBiT-G protein dissociation assay and the NanoBiT-β-arrestin recruitment assay of C286^6.47^A and H291^6.52^A mutants. Dashed lines in the mutant panels represent the wild-type (WT) KOR (1:1) response. Data are presented as mean values ± SEM (n = 3). Note that in numerous data points, the error bars are smaller than the size of the symbols, making them not visible. (d) Interaction between C286^6.47^ and T321^7.43^ and conformational changes in TM6 during agonists and antagonist binding. Arrows indicate the extent of conformational change in TM6 when comparing inactive and active forms. (e) Interaction network formed between agonists, H291^6.52^ and N322^7.44^, which links with a toggle switch (W287^6.48^) that is common to GPCRs and important for their activation. (f) Interaction of H291^6.52^ and W287^6.48^ in agonist- or antagonist-bound states. Nalfurafine-bound KORs are represented in gray, U-50,488H-bound KORs is in orange, and inverse agonist JDTic-bound KORs are displayed in magenta.

### Structural comparison between the nanobody-bound KOR and the KOR-G_i_ complex in nalfurafine-bound state

Recently, the structure of the nalfurafine-bound KOR was elucidated using X-ray crystallography in its nanobody (Nb39)-bound state, which stabilizes its active conformation^14^. In this arrangement, both the C-terminal regions of Nb39 and G_i_ enter and bind to the intracellular pocket of the KOR.

A comparative structural analysis of the KOR complex in these nalfurafine-bound states highlighted minor conformational variations in amino acid side chains, along with three major differences. Firstly, in the Nb39-bound state, we observed a notable outward opening of the intracellular region of TM5 and TM6, as illustrated in Supplementary Figure 12a. Although the C-terminal region of G_i_ penetrates the KOR more deeply compared to Nb39, the loop region of Nb39 exhibits significant interaction with the KOR pocket, likely influencing the observed conformational changes in TM5 and TM6 (Supplementary Fig. 12b). Secondly, a noticeable difference was observed in the orientation of the side chains of residues Y140^3.34^ and W183^4.50^ (Supplemental Fig. 12c). In the nalfurafine-bound form of the KOR-G_i_ complex, the side chain of W183^4.50^ underwent a significant shift, resulting in the formation of a hydrogen bond with Y140^3.34^. To evaluate the importance of the interaction between Y140^3.34^ and W183^4.50^ on G-protein dissociation and arrestin-recruitment, we examined the signaling activity of Y140^3.34^F and W183^4.50^A mutants. Surprisingly, these amino acid substitutions did not substantially affect G-protein dissociation and arrestin-recruitment activities. Furthermore, in the presence of U-50,488H, the conformations of Y140^3.34^ and W183^4.50^ closely resembled those observed in the nalfurafine-bound KOR-Nb39 structure (Supplementary Fig. 12c). Additionally, slight differences in the rotamers of amino acid side-chains involved in arrestin recruitment activity were found in this study (Supplementary Fig. 12d).

## DISCUSSION

The aim of this investigation was to leverage structural biology, infrared spectroscopy, and pharmacological analysis to pinpoint crucial amino acid residues governing the signaling activity of KOR and to unravel the molecular basis of biased agonism in GPCRs. Alongside the static structural data of G protein-bound KOR acquired with cryo-EM, insights into the conformational changes of KORs preceding G protein binding were gleaned from dynamic infrared spectroscopy, enabling the identification of amino acids involved in signal transduction. K227^5.40^ and Y312^7.34^ emerged as pivotal residues for arrestin-recruitment, as evidenced by cryo-EM data and pharmacological analysis, whereas C286^6.47^ and H291^6.52^ were identified through FT-IR and pharmacological analysis (Fig. 5). This investigation yields promising insights into these amino acids from a drug discovery standpoint.

**Figure 5:**
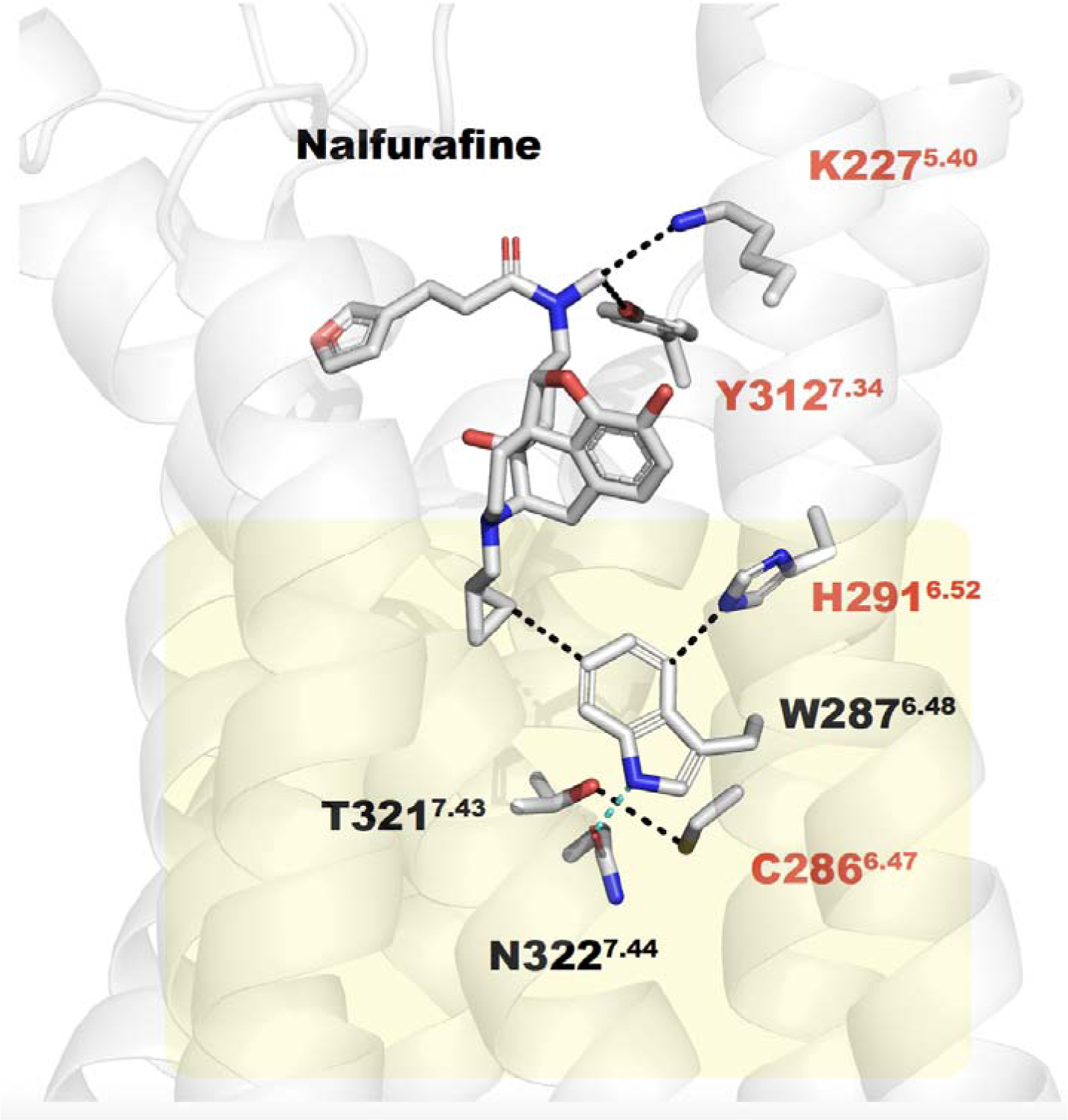
Amino acid residues within TM5, 6, and 7 that are involved in arrestin recruitment in KOR. K227^5.40^ and Y312^7.34^ interact directly with the agonist, facilitating signal transducer-specific binding. On the contrary, alterations within the network of amino acid residues between TM6 and TM7 (highlighted in yellow) influence the selective binding of signaling factors. Amino acid residues crucial for arrestin recruitment, as identified in this study, are denoted by red letters.

Previous studies have implicated two amino acids, K227^5.40^ and Y312^7.34^, in the signaling properties of KOR^14,15^. This study adds further clarity to their roles in signaling selectivity. The side chain of K227^5.40^ showed weak ion binding to the side chain of E297^6.58^ in the U-50,488H-bound KOR structure, while in the nalfurafine-bound KOR structure, it interacted with the agonist. The K227^5.40^A mutant exhibited a significant decrease in arrestin-recruitment activity upon the addition of agonists, and G protein-coupling activity was slightly worse in U-50,488H-bound form. From a drug discovery standpoint, we propose that agonists that strongly interact with K227^5.40^ and inhibit the interaction between K227^5.40^ and E297^6.58^ are G protein-biased agonists.

Y312^7.34^ also provided important insights into the development of biased ligands. Specifically, the Y312^7.34^F mutant reduced arrestin-recruitment activity while maintaining predominant G protein-coupling activity. Designing compounds with weaker affinities for the side chain of Y312^7.34^ than nalfurafine is expected to reduce arrestin-recruitment activity while preserving G protein-coupling activity. Che *et al.* reported the importance of Y312^7.34^ in biased activity, as they observed biased activity with biased ligands of MOR by replacing Y312^7.34^ of KOR with a tryptophan residue. In this study, we elucidated the structures of two KOR-selective agonist binding states and conducted a detailed structural and pharmacological comparison. We found that the interaction of the ligand with Y312^7.34^ is pivotal for biased activity. Zhuang *et al.* determined the structures of several biased/balanced agonist-bound MOR-G protein complexes and suggested the importance of reducing interaction with TM6/7 in the design of G protein-biased agonists^20^. This proposal for MOR contradicts the findings of our study, where it is crucial for G protein-biased agonists to interact appropriately with Y312^7.34^ at TM7 of KOR. Therefore, to develop analgesics without the side effects of KOR, it is imperative to design agents that differ from those targeting MOR.

In contrast, the involvement of C286^6.47^ and H291^6.52^ of TM6 in signaling selectivity, as discovered through FT-IR measurements and mutant analysis in this study, has not been previously documented. Structural analysis and FT-IR results from our study suggest that these amino acids do not directly interact with the agonist but rather modify the interaction network with other amino acids in TM7 upon agonist binding.

Furthermore, the alanine mutants exhibit diminished arrestin-recruitment activity, indicating the significance of the interaction between these amino acids in TM6 and those in TM7 for biased signaling (Fig. 5). Specifically, the interaction between C286^6.47^ and T321^7.43^ is crucial for arrestin recruitment while not affecting G protein-coupling activity. Hence, the design of agonists or allosteric ligands that disrupt these interactions could potentially lead to the development of KOR agonists devoid of side effects.

H291^6.52^ forms an intricate interaction network with T322^7.44^ via W287^6.47^, and the reduced arrestin-recruitment activity observed in the H291^6.52^A mutant may stem from the disruption of these networks. In essence, the H291^6.52^A mutation could alter the orientation of the side chain of W287^6.47^, consequently affecting the interaction between W287^6.47^ and N322^7.44^ of TM7. In summary, this study successfully identifies the amino acid residues crucial for arrestin recruitment as well as the amino acid residues that interact with agonists or whose interaction networks are altered by agonist binding (Fig. 5).

Our integrated approach, amalgamating structural analysis and infrared spectroscopy, elucidates a pivotal signaling mechanism crucial for drug discovery. This methodology, directly applicable to opioid receptors like KORs, holds promise for extension to other GPCRs. Approaches akin to this study show potential in facilitating the rational design and enhancement of therapeutics, encompassing biased agonists such as nalfurafine for KORs, and can be extrapolated to the development of biased agonists for numerous other GPCRs.

## METHODS

### Cells

*Spodoptera frugiperda* 9 (Sf9) insect cells were cultured in ESF-921 (Wako), 50 units/mL penicillin, 50 μg/mL streptomycin (Wako), and 0.5 μg/mL amphotericin B at 27 °C. Parental human embryonic kidney 293 (HEK293) cells^21^ were grown in Dulbecco’s modified Eagle’s medium (DMEM; Nacalai Tesque) supplemented with 10% FBS (Nichirei Biosciences), 100 units/mL penicillin, and 100 μg/mL streptomycin (Nacalai Tesque) at 37 °C in a 5% CO_2_ incubator.

### Constructs

For the human KOR construct, the N-terminal (1-53) and C-terminal (359-380) regions were deleted from the wild-type KOR based on the previously reported crystal structure of nanobody-stabilized KOR^14,15^, and the deleted N-terminal (1-53) was replaced with b562RIL (BRIL). An N-terminal A 3C protease site was inserted between BRIL and KOR, and a 3C protease site was also inserted at the C-terminus of KOR to introduce GFP and 10xHis tag. The sf9 expression vector for the heterotrimeric G protein was provided by Brian K Kobilka’s lab, with G_i_ introduced into pFastBac and G_β_ and G_γ_ into the pFastBac Dual vector. Sf9 expression vector with scFv16 introduced into the pFastBac vector was provided by Nureki Lab^22^.

### Expression and purification of the human KOR-G_i1_Gβ_1_γ_2_-scFv16 complex

Recombinant baculoviruses were produced using the Bac-to-Bac system (Invitrogen) and infected Spodoptera frugiperda (Sf9) cells. Briefly, Sf9 cells were cultured in ESF921 medium at a density of 3-4 x 10^6^ cells/ml and co-infected with three different baculoviruses: KOR, G_i1_, and G_β1γ2_ in a 1:2:1 ratio. 27 °C, 125 rpm, 48-72 hours. After infection for 2 hours, cells were collected by centrifugation, and cell pellets were stored at −80 °C. Insect cell membranes were resuspended in suspension buffer containing 30 mM HEPES-NaOH (pH 7.5), 750 mM NaCl, 5 mM imidazole, 1 mg/ml iodoacetamide, 1 µM agonist, 10 mM leupeptin, 1 mM benzamidine, and apyrase. The suspension was incubated at 4 °C for 30 min; DDM and CHS were added at final concentrations of 1% (w/v) and 0.2% (w/v), respectively, and incubation continued for another 2 hr. The supernatant was ultracentrifuged at 100,000 g for 30 min and purified with Ni-NTA Superflow resin (Qiagen). The purified supernatant was applied to a column (GFPNb column) conjugated with a GFP recognition antibody (GFP nanobody) and washed with 3 volumes of buffer (30 mM HEPES-NaOH (pH 7.5), 750 mM NaCl, 0.1% (w/v) DDM, 0.03% (w/v) CHS, 10 µM (agonist) and washed. DDM was then gradually replaced over 1 hr with buffer containing 0.01% (w/v) lauryl maltose neopentyl glycol (MNG), and the NaCl concentration was lowered to 100 mM. KOR-G_i_G_bg_ complex was eluted by adding 3C protease in the presence of 10 µM agonist. The preparation of the GPCR-G protein complex using the GFPnanobody column was similar to our previously reported method for the prostaglandin receptor-G_i_G_bg_ complex^22^. scFv16 was purified as in the previously reported study of the structural analysis of the MT1- G_i_G_bg_-scFv16 complex^22^. The purified KOR-G_i_G_bg_ complex and scFv16 were combined in a 1:2 molar ratio and incubated overnight at 4 °C. The resulting KOR-G_i_G-scFv16 complex was purified using a Superdex 200 Increase 10/300 GL column (Cytiva) in 20 mM HEPES-NaOH (pH 7.5), 100 mM NaCl, 0.001% (w/v) MNG, 0.0003% GDN, 0.001 % (w/v) CHS, 1 μM agonist, and further purified using size exclusion chromatography in a buffer of 100 mM TCEP (Supplemental Figs. 1a-d). The eluted peak fraction was concentrated to approximately 5 mg/ml using an Amicon Ultra concentrator (Millipore) with a molecular weight cutoff of 100 kDa.

### Cryo-EM grid preparation and data collection

A 3 µl aliquot of sample solution was applied to a glow discharge QUANTIFOIL R1.2/1.3 Au 300 mesh grid (Quantifoil Micro Tools GmbH) on a Vitrobot Mark IV (Thermo Fisher Scientific). After wiping off the excess solution on the grid with filter paper, the samples were rapidly frozen in liquid ethane. The frozen grids were screened using a Talos Glacios cryo-transmission electron microscope (cryo-TEM) (operated at 200 keV and equipped with a Falcon 4 direct electron detector (Thermo Fisher Scientific)) at Institute for Life and Medical Sciences, Kyoto University, to check sample conditions such as ice thickness and particle dispersion. Cryo-EM data collection was performed using a Titan Krios cryo-TEM (Thermo Fisher Scientific) in EFTEM nanoprobe mode operating at 300 keV equipped with a Cs corrector (CEOS GmbH) at the Institute for Protein Research, Osaka University. Images were acquired as movies using a Gatan BioQuantum energy filter with a slit width of 20 eV and a K3 direct detection camera (Gatan, Inc). For U-50,488H or nalfurafine-bound KOR-G_i_ complexes, a total of 18,885 and 14,349 movies were collected, respectively, at a pixel size of 0.675 Å^2^ and a total dose of 60 e^-^/Å^2^ Automated data collection was performed using a 3×3 hole-pattern beam-image-shift scheme with a nominal defocus range of −0.7 to −1.5 μm facilitated by the SerialEM software^24^.

### Image processing

Image processing was performed with cryoSPARC^25^. Micrographs were motion-corrected using patch motion correction, and CTF parameters were estimated using patch CTF estimation. For U-50,488H- or nalfurafine-bound KOR datasets, particles were initially picked using a blob picker (minimum particle size 100, maximum particle size 200). Subsequently, particle extraction was performed with a particle box size of 360 pixels, resulting in the extraction of 4,148,764 particles from the U-50,488H-bound KOR dataset and 7,066,947 particles from the nalfurafine-bound KOR dataset. Only in the case of U-50,488H-bound KOR, once 2D classification was done, Topaz extraction was performed, and 4,148,764 particles were extracted from 18,845 micrographs. After multiple rounds of 2D classification, we selected 3,078,397 or 3,957,130 particles for *ab-initio* reconstruction. In the case of the data of the nalfurafine-bound KOR, a set of 2,439,864 particles was further classified into two classes using ab-initio reconstruction followed by heterogenous refinement. Particles from one class (1,225,096 or 858,423 particles) were combined and subjected to homogeneous refinement and non-uniform refinement, yielding a resolution of 2.90 Å and 2.76 Å for U-50,488H- and nalfurafine-bound KOR-G_i_ complex, respectively. Resolution is calculated using the “gold standard” Fourier Shell Correlation (FSC) between two independently refined maps reconstructed independently from randomly divided half datasets. The image processing steps are summarized in Supplemental Figs 2 and 3.

### Model building and refinement

The atomic model building was performed by manual iterative building in Coot^26^, followed by refinement with phenix.real_space_refine in the Phenix program suite^27^. In this model, 97% of the residues were in the favored regions of the Ramachandran plot, and all the others were in the allowed regions. Refinement statistics are shown in Supplementary Table 2.

### Preparation of proteoliposome of KOR

For ATR-FTIR spectroscopy measurements, KOR solubilized in detergent was reconstituted into asolectin liposomes with a 10-fold molar excess. Surfactant removal was achieved by incubating with Biobeads SM-2 (Bio-Rad). After removing the Biobeads, the reconstituted KOR was recovered by ultracentrifugation. Following several cycles of washing/spinning, the lipid-reconstituted KOR was suspended in a buffer consisting of 5 mM phosphate (pH 7.5) and 10 mM KCl. Furthermore, the determination of detergent removal was based on the infrared absorption peaks attributed to detergent (observed in the 1200 cm^-1^ region) in the ATR-FTIR measurements.

### Ligand binding-induced ATR-FTIR difference spectroscopy

A 3 μL suspension of lipid-reconstituted KOR was dropped onto the surface of a silicon ATR crystal (with 3 internal reflections). After gently drying to form a thin film sample by natural drying, the film sample was rehydrated through a flow cell maintained at 20 °C by circulating water at a flow rate of 0.4 mL min-^1^, with a solvent containing 200 mM phosphate buffer (pH 7.5) containing 140 mM KCl and 3 mM MgCl_2_.

Before measurements, 0.1 mM naltrexone was perfused into the buffer to bind KOR with Naltrexone. Using an FTIR spectrometer equipped with a liquid nitrogen-cooled MCT detector (Bio-Rad FTS6000, Agilent, CA, USA), ATR-FTIR absolute absorbance spectrum of the inactive form of KOR bound to Naltrexone was first recorded at a resolution of 2 cm^-1^ (averaging 64 spectra). Next, ATR-FTIR absolute absorbance spectrum was recorded in a second buffer solution containing 0.01 mM Nalfurafine or U-50,488 added, during which ligands binding to KOR were exchanged.

The ATR-FTIR difference spectrum was obtained by subtracting the absolute absorbance spectrum of the Naltrexone-bound form from the absolute absorbance spectrum of the Nalfurafine or U-50,488-bound form. This cycling procedure was repeated 3 to 6 times. Corrections were made for protein/lipid contraction, water vapor, CO_2_, and contributions from water/buffer components in the spectra.

### Plasmids

For the NanoBiT-G-protein dissociation assay, the full-length human KOR was inserted into the pCAGSS expression vector with the N-terminal fusion of the hemagglutinin-derived signal sequence (ssHA), FLAG epitope tag and a flexible linker (MKTIIALSYIFCLVFA<u>DYKDDDDK</u>GGSGGGGSGGSSSGGG; the FLAG epitope tag is underlined). The resulting construct was named ssHA-FLAG-KOR. For the NanoBiT-β-arrestin-recruitment assay, the ssHA-FLAG-KOR construct was C-terminally fused with the flexible linker, and the SmBiT fragment GGSGGGGSGGSSSGG<u>VTGYRLFEEIL</u>; the SmBiT is underlined). The resulting construct was named ssHA-FLAG-KOR-SmBiT.

### NanoBiT-G protein-dissociation assay

KOR ligand-induced G protein dissociation was measured by the NanoBiT-G-protein dissociation assay^28^, in which the interaction between a Gα subunit and a Gβγ subunit was monitored by the NanoBiT system (Promega). Specifically, a NanoBiT-G_i_ protein consisting of the Gα_i1_ subunit fused with a large fragment (LgBiT) at the α-helical domain (between the residues 91 and 92 of Gα_i1_; Gα_i1_-LgBiT) and the N-terminally small fragment (SmBiT)-fused Gγ_2_ subunit with a C68S mutation (SmBiT-Gγ_2_-CS) was expressed along with untagged Gβ_1_ subunit. HEK293A cells (Thermo Fisher Scientific, cat no. R70507) were seeded in a 6-well culture plate at a concentration of 2 x 105 cells ml^-1^ (2 ml per well in DMEM (Nissui) supplemented with 5 % fetal bovine serum (Gibco), glutamine, penicillin, and streptomycin), one day before transfection. Transfection solution was prepared by combining 6 µL (per dish hereafter) of polyethylenimine (PEI) Max solution (1 mg ml^-1^; Polysciences), 200 µL of Opti-MEM (Thermo Fisher Scientific), and a plasmid mixture consisting of 200 ng ssHA-FLAG-KOR (or an empty plasmid for mock transfection), 100 ng Gα_i1_-LgBiT, 500 ng Gβ_1_ subunit and 500 ng SmBiT-Gγ_2_-CS subunit. To compare expression-matched KOR mutants, the WT KOR plasmid was serially titrated. After incubation for 1 day, the transfected cells were harvested with 0.5 mM EDTA-containing Dulbecco’s PBS, centrifuged, and suspended in 2 ml of HBSS containing 0.01 % bovine serum albumin (BSA; fatty acid-free grade; SERVA) and 5 mM HEPES (pH 7.4) (assay buffer). The cell suspension was dispensed in a white 96-well plate at a volume of 80 µL per well and loaded with 20 µL of 50 µM coelenterazine (Angene) diluted in the assay buffer. After a 2 hr incubation at room temperature, the plate was measured for baseline luminescence (SpectraMax L, Molecular Devices), and titrated concentrations of a KOR ligand (nalfurafine or U-50,488H; 20 µL; 6X of final concentrations) were manually added. The plate was immediately read for the second measurement as a kinetics mode and luminescence counts recorded from 5 to 10 min after compound addition were averaged and normalized to the initial counts. The fold-change values were further normalized to those of vesicle-treated samples and used to plot the G-protein dissociation response. Using the Prism 9 software (GraphPad Prism), the G-protein dissociation signals were fitted to a four-parameter sigmoidal concentration-response curve with a constraint of the *HillSlope* to absolute values less than 2. For each replicate experiment, the parameters *Span* (= *Top* – *Bottom*), pEC_50_ (negative logarithmic values of EC_50_ values), and *Span*/EC_50_ of the individual KOR mutants were normalized to those of WT KOR performed in parallel, and the resulting parameters *E_max_*, ΔpEC_50_ and relative intrinsic activity (RAi), respectively, were used to denote ligand response activity of the mutants.

### NanoBiT-**β**-arrestin-recruitment assay

Ligand-induced β-arrestin-recruitment assay was measured as described previously ^29^. Transfection was performed according to the same procedure as described in the “NanoBiT-G protein-dissociation assay” section, except for a plasmid mixture consisting of 500 ng ssHA-FLAG-KOR-SmBiT and 100 ng N-terminally LgBiT-fused β-arrestin2. The transfected cells were dispensed into a 96-well plate, and ligand-induced luminescent changes were measured by following the same procedures as described for the NanoBiT-G protein-dissociation assay.

### Flow cytometry

Transfection was performed according to the same procedure as described in the “NanoBiT-G protein-dissociation assay” and the “NanoBiT-β-arrestin-recruitment assay” sections. One day after transfection, the cells were collected by adding 200 μl of 0.53 mM EDTA-containing Dulbecco’s PBS (D-PBS), followed by 200 μl of 5 mM HEPES (pH 7.4)-containing Hank’s Balanced Salt Solution (HBSS). The cell suspension was transferred to a 96-well V-bottom plate in duplicate and fluorescently labeled with an anti-FLAG epitope (DYKDDDDK) tag monoclonal antibody (Clone 1E6, FujiFilm Wako Pure Chemicals; 10 μg per ml diluted in 2% goat serum- and 2 mM

EDTA-containing D-PBS (blocking buffer)) and a goat anti-mouse IgG secondary antibody conjugated with Alexa Fluor 488 (Thermo Fisher Scientific, 10 μg per ml diluted in the blocking buffer). After washing with D-PBS, the cells were resuspended in 200 μl of 2 mM EDTA-containing-D-PBS and filtered through a 40 μm filter. The fluorescent intensity of single cells was quantified by an EC800 flow cytometer equipped with a 488 nm laser (Sony). The fluorescent signal derived from Alexa Fluor 488 was recorded in an FL1 channel, and the flow cytometry data were analyzed with the FlowJo software (FlowJo). Live cells were gated with a forward scatter (FS-Peak-Lin) cutoff at the 390 setting, with a gain value of 1.7. Values of mean fluorescence intensity (MFI) from approximately 20,000 cells per sample were used for analysis. For each experiment, we normalized an MFI value of the mutants by that of WT performed in parallel and denoted relative levels.

## QUANTIFICATION AND STATISTICAL ANALYSIS

In the signaling assay, maximum luminescence intensity post stimulation was quantified. The luminescence intensity reached a plateau about 10 min after stimulation. Each point represents the mean value ± s.e.m. All the measurements were performed in triplicate. Sigmoid curve fitting was performed with Prism 7 (GraphPad).

### Lead Contact

Further information and requests for resources and reagents should be directed to and will be fulfilled by the lead contact, Ryoji Suno and Takuya Kobayashi.

### Data and Code Availability

The cryo-EM reconstructions generated in this study have been deposited into the Electron Microscopy Data Bank (EMDB) with accession numbers EMD-37545 and 38751. The 3D models reported in this paper have been deposited in the Protein Data Bank with accession code PDB ID: 8WHR and 8XXJ.

## ACKNOWLEDGEMENTS

The plasmids for expressing scFv16 were provided by Hiroyuki Okamoto (Tokyo University). This research was supported by the Takeda Science Foundation (to R.S., Y.S., and T. Kobayashi), AMED Core Research for Evolutional Science and Technology (CREST) (JP21gm0910007 to T. Kobayashi), AMED Science and Technology Platform Program for Advanced Biological Medicine under Grant Number JP21am0401020 (to T. Kobayashi), AMED Research on Development of New Drugs (JP20ak0101103 to T. Kobayashi), The Naito Foundation (to T. Kobayashi), Koyanagi Foundation (to T. Kobayashi.), FOREST Program JPMJFR215T (to Asuka Inoue) and JST Moonshot Research and Development Program JPMJMS2023 (to Asuka Inoue) from Japan Science and Technology Agency (JST), JSPS KAKENHI (19H03428 to R. Suno.; 21H04791, 21H05113 and JPJSBP120218801 to Asuka Inoue), Grant-in-Aid for Transformative Research Areas (21H05111 to R. Suno., Asuka Inoue, and T.S., 21H05112 to R. Suno, 21H05113 to Asuka Inoue, and 21H05115 to T.S.) and for MEXT LEADER Program (to Y.S.), and Grant-in-Aid for Specially Promoted Research (JP21H04969 to H.K). The authors thank Kayo Sato, Shigeko Nakano, and Ayumi Inoue at Tohoku University for their assistance in plasmid preparation. Cryo-EM analysis was supported by the Cooperative Research Program (Joint Usage/Research Center program) of Institute for Life and Medical Sciences, Kyoto University, and the Platform Project for Supporting Drug Discovery and Life Science Research (Basis for Supporting Innovative Drug Discovery and Life Science Research (BINDS)) from AMED (JP23ama121001 to T. Kato).

## Author Contributions

C.S. carried out protein expression and purification of the receptor, G proteins, GFP nanobody, and scFv16 fragment. T.S. synthesized the compounds. C.S., T.T., Akitoshi Inoue, E.A created a mutant construct of KOR. C.S. prepared the cryo-EM sample of the KOR-Gi-scFv16 complex. C.S., R. Suno, Y.S., and M.H. carried out the cryo-EM data collection. R. Suno carried out cryo-EM data processing and model building of the KOR-Gi-scFv16 complex. R. Suzuki and T.O. carried out the signaling assays. R.N. and S.I. performed the infrared spectroscopy measurements. R. Suno designed the project, and H.K., K.K., T. Kato. and T. Kobayashi supervised the overall project. C.S. and R. Suno wrote the manuscript. All authors discussed the results and commented on the manuscript.

