## Supplementary Information for "Structural and Dynamic Insights into the Biased Signaling Mechanism of the Human Kappa Opioid Receptor"

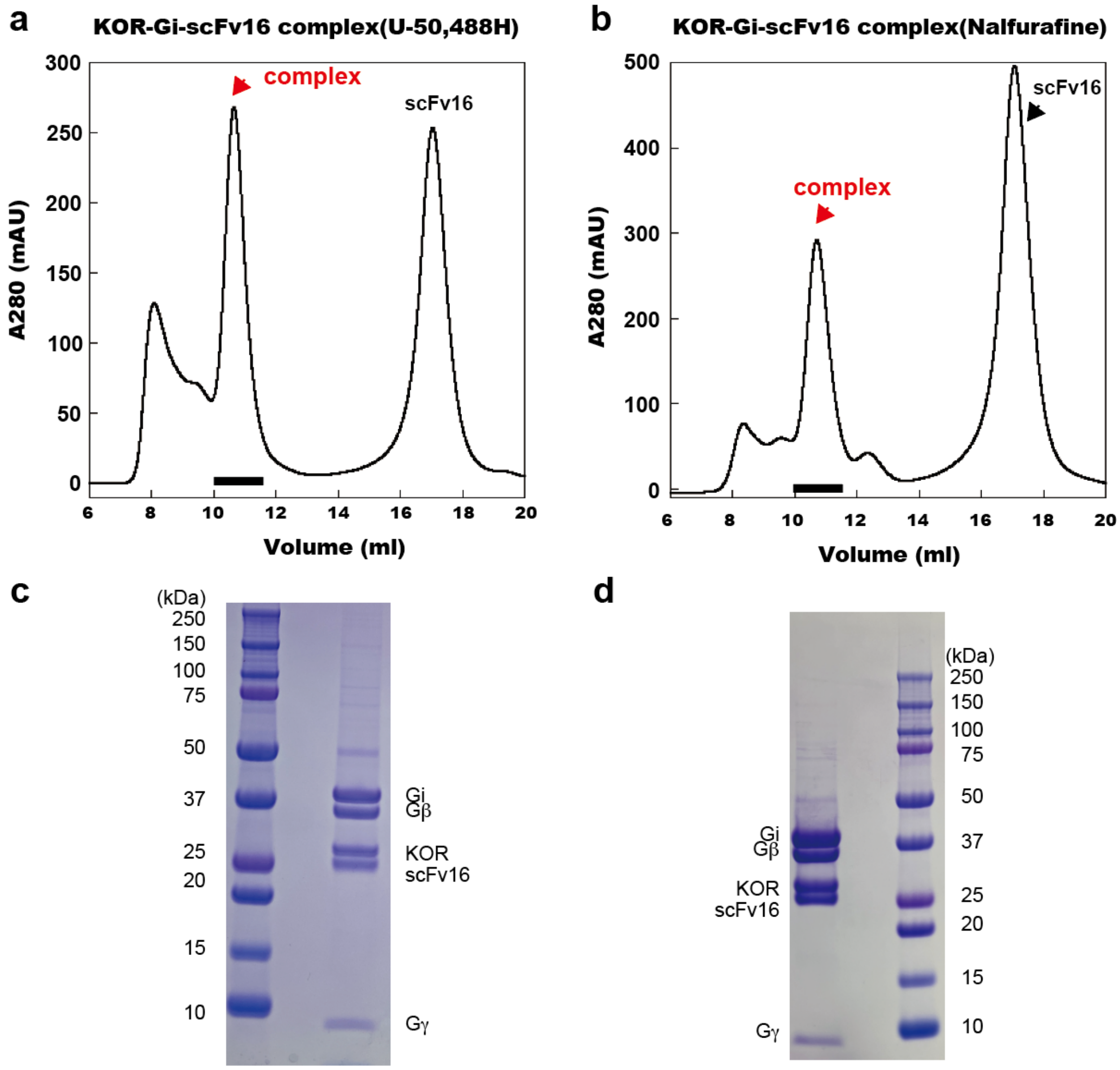


**Supplemental Figure 1.** Preparation of KOR-$G_i$ signaling complexes in U-50,488H-(a,c) and nalfurafine (b,d)-bound states. Chromatogram of gel filtration chromatography purification (a, b). SDS-PAGE of complex peaks in gel filtration chromatography (c, d).

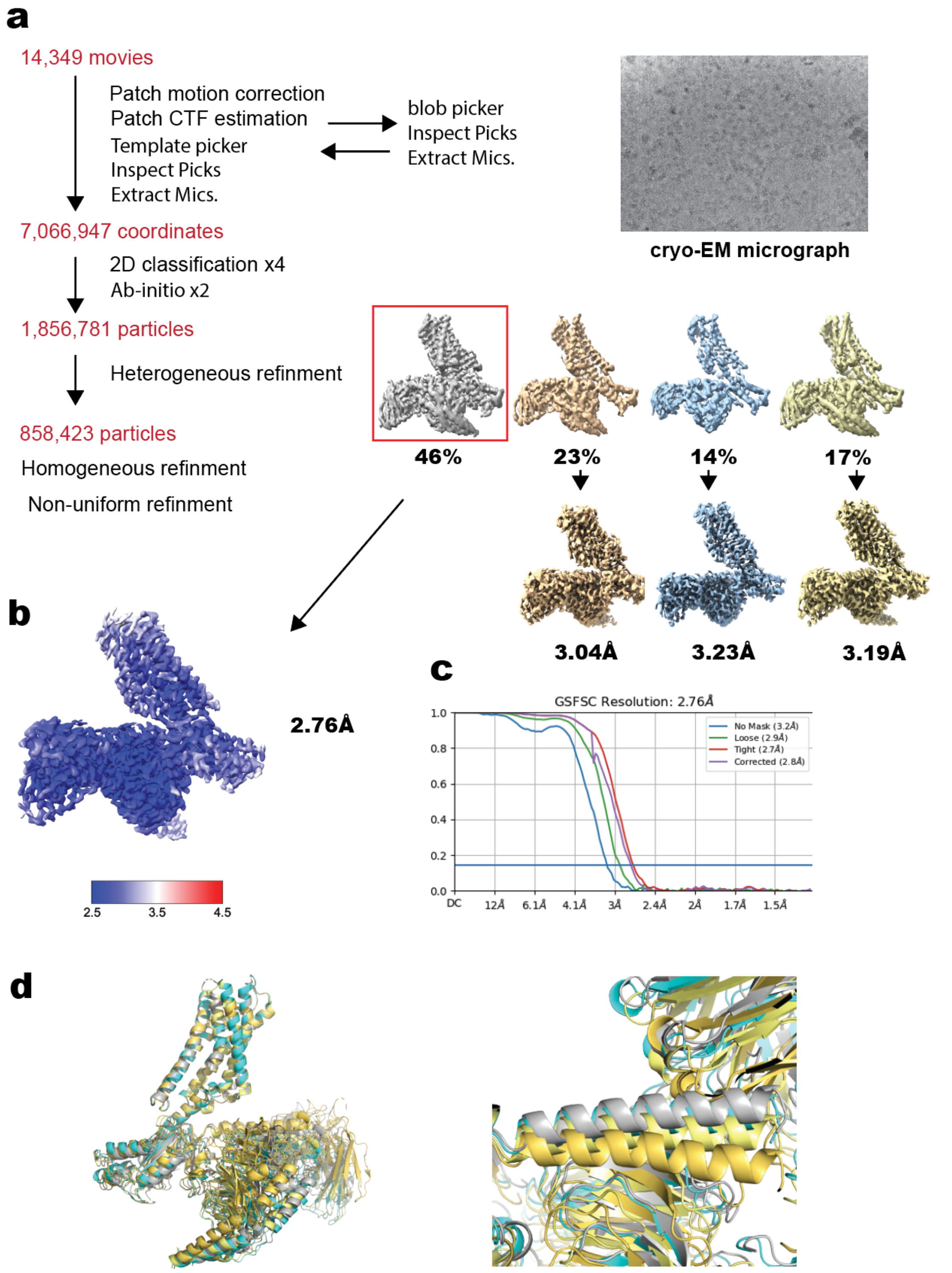


**Supplemental Figure 2. Cryo-EM data processing of nalfurafine-bound KOR-Gi complex**

a. Workflow of the cryo-EM data processing. **b.** Local resolution maps for the non-uniform refinement of nalfurafine-bound KOR-$G_i$ signaling complex. **c.** Gold standard

FSC plots for the non-uniform refinement of nalfurafine-bound KOR-$G_i$ signaling complex calculated in cryoSPARC. d. Superimposed view of the KOR-$G_i$ signaling complex in four different nalfurafine binding states. Relative differences in G protein position (left) and N-terminal orientation (right) of G proteins due to superposition of receptor regions. The four KOR-$G_i$ signaling complex structures are at 2.76, 3.04, 3.23, and 3.19 Å resolution, respectively, and are shown in gray, orange, cyan, and yellow, respectively.

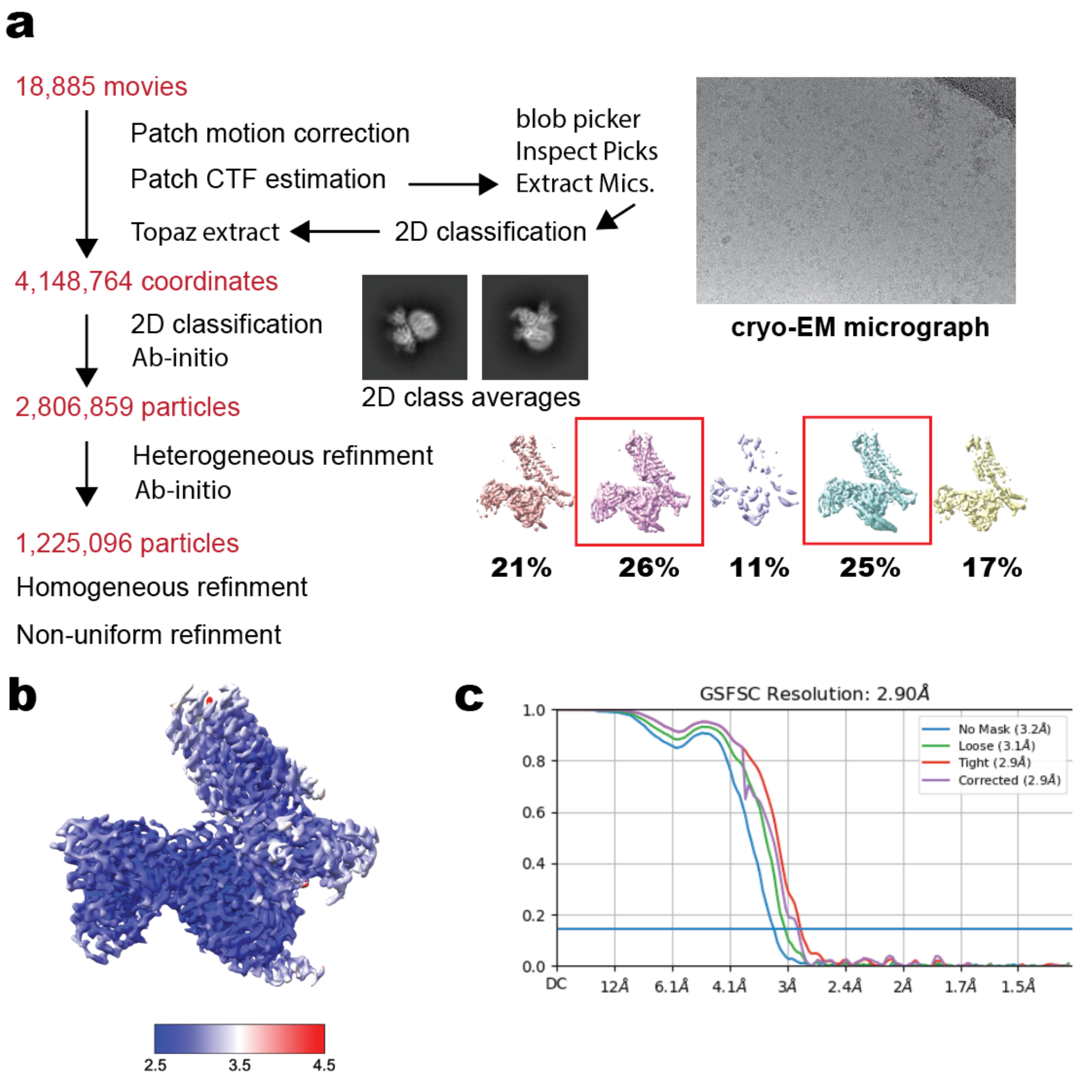


**Supplemental Figure 3. Cryo-EM data processing of U-50,488H-bound KOR-$G_i$ complex**

**a.** Workflow of the cryo-EM data processing. *Ab initio* reconstruction was performed by mixing the particles in the two 3D classes indicated by red squares from the heterogeneous refinement results. **b.** Local resolution maps for the non-uniform refinement of U-50,488H-bound KOR-$G_i$ signaling complex. **c.** Gold standard FSC plots for the non-uniform refinement of U-50,488H-bound KOR-$G_i$ signaling complex calculated in cryoSPARC.

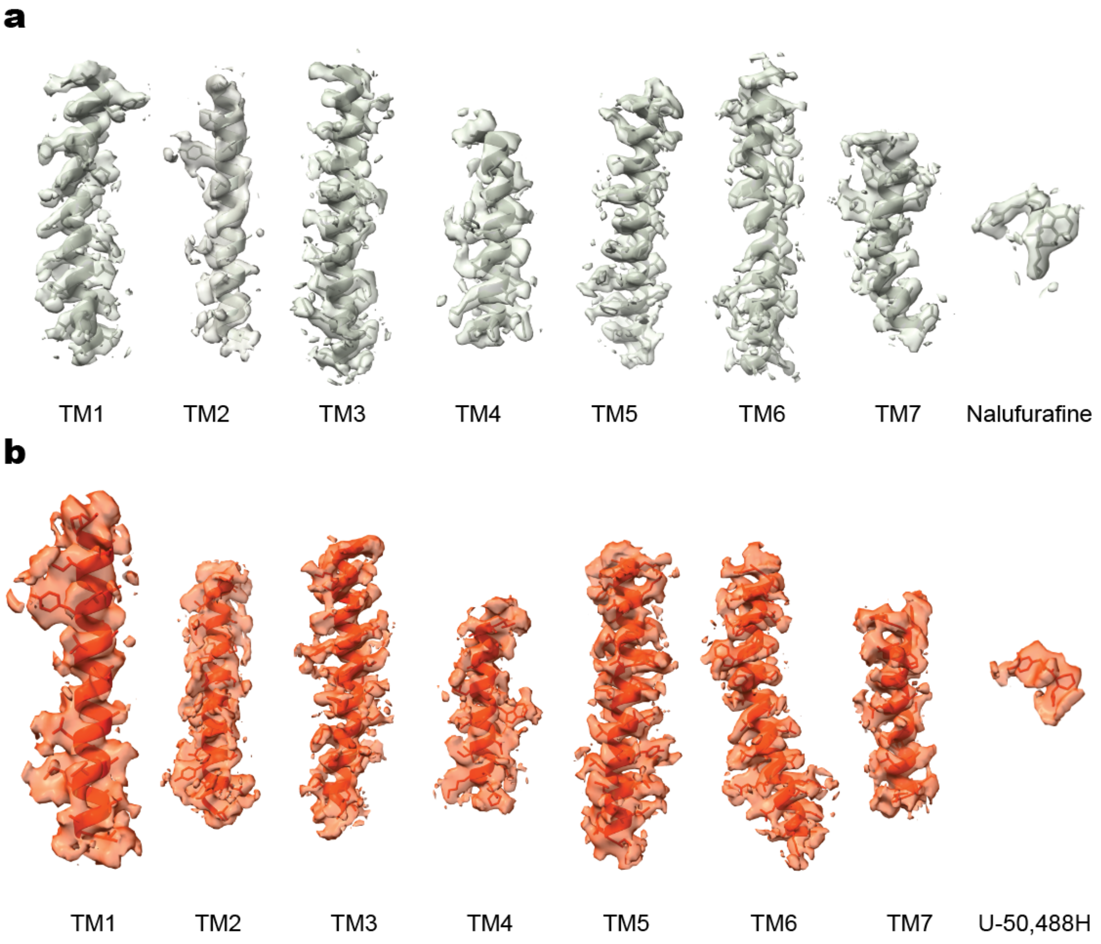


**Supplemental Figure 4. Cryo-EM density maps and models of the seven transmembrane helices (TM1-7) of nalfurafine- (a) or U-50,488H- (b) bound KOR.** Maps are shown in gray and orange, respectively.

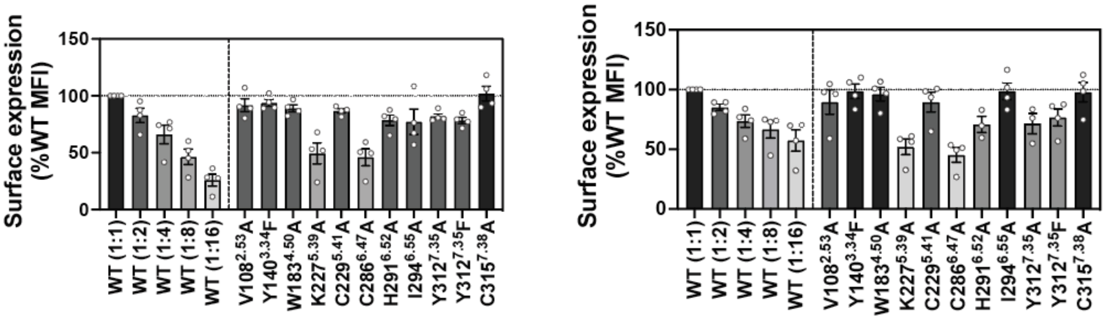


**Supplemental Figure 5. Surface expression of the KOR mutants analyzed by flow cytometry.**

Data are presented as mean values ± SEM ($n$ = 3-4; dots). The left is the results for the constructs used for G-protein dissociation and the right is those for the constructs used for arrestin recruitment, which are fused with Sm-BiT in its C terminus. The dilutions in the WT indicate volumes of the transfected WT KOR plasmid. Mutants with lowered surface expression levels are shown as grey bars corresponding to relative receptor expression levels.

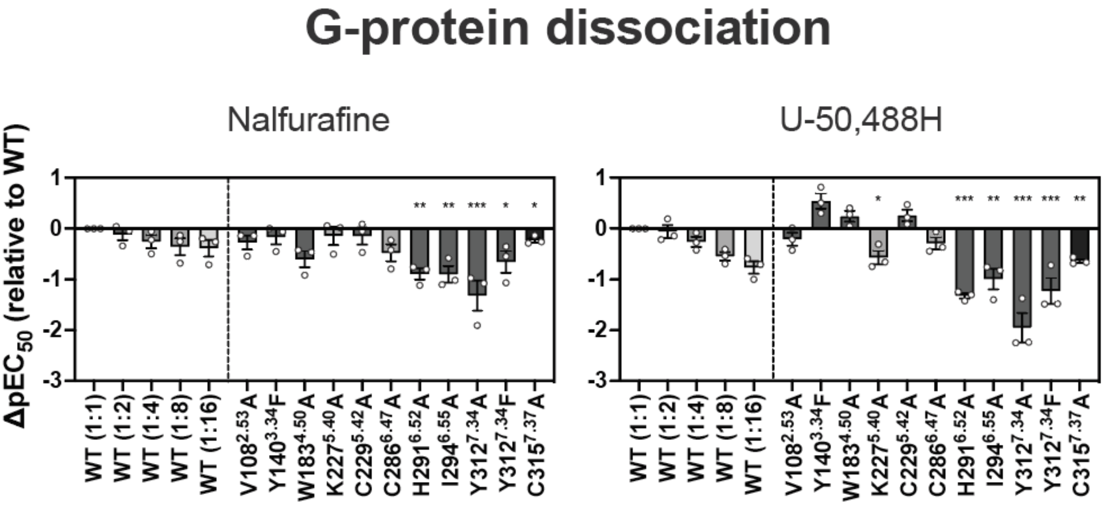


**Supplemental Figure 6.** **$G_i$-coupling activity analyzed by the NanoBiT-G protein dissociation assay.**

Pharmacological parameters for the $G_i$-coupling activity analyzed by the NanoBiT-G-protein dissociation assay. Data are presented as mean values ± SEM (*n* = 3-4; dots). For the individual experiments performed in parallel, data were normalized to the wild-type (WT) KOR (1:1) and presented as $\Delta pEC_{50}$. The colors in the mutant bars correspond to the expression-matched WT conditions. Statistical analyses were performed using *t*-tests and the ordinary one-way ANOVA followed by Dunnett test with the expression-matched (colored) WT response. ns, $p > 0.05$; $*p < 0.05$; $**p < 0.01$; $***p < 0.001$. Data are presented as mean values ± SEM (*n*=3; dots).

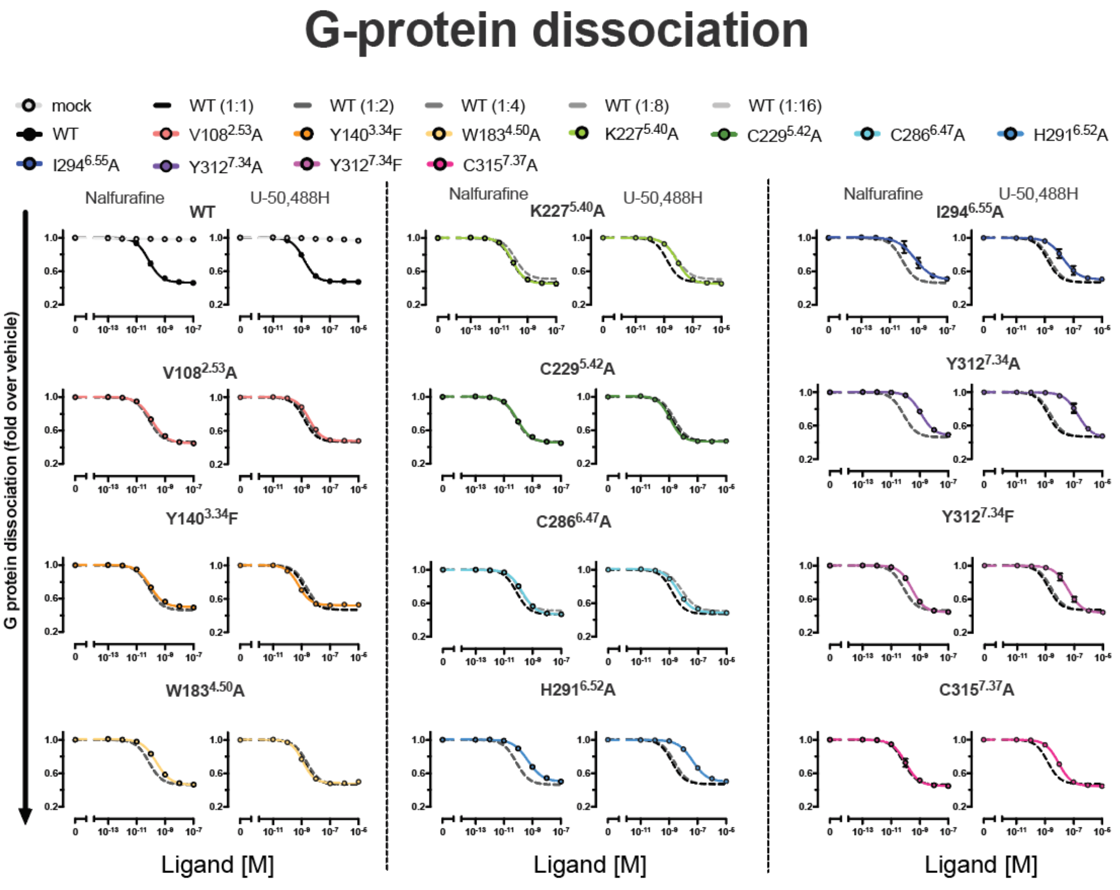


**Supplemental Figure 7. Concentration-response curves of the NanoBiT-G protein dissociation assay.**

Dashed lines in the mutant panels represent the wild-type (WT) KOR (1:1, 1:2, 1:4, 1:8, or 1:16) response. Data are presented as mean values ± SEM (*n* = 3). Note that in numerous data points, the error bars are smaller than the size of the symbols, making them not visible.

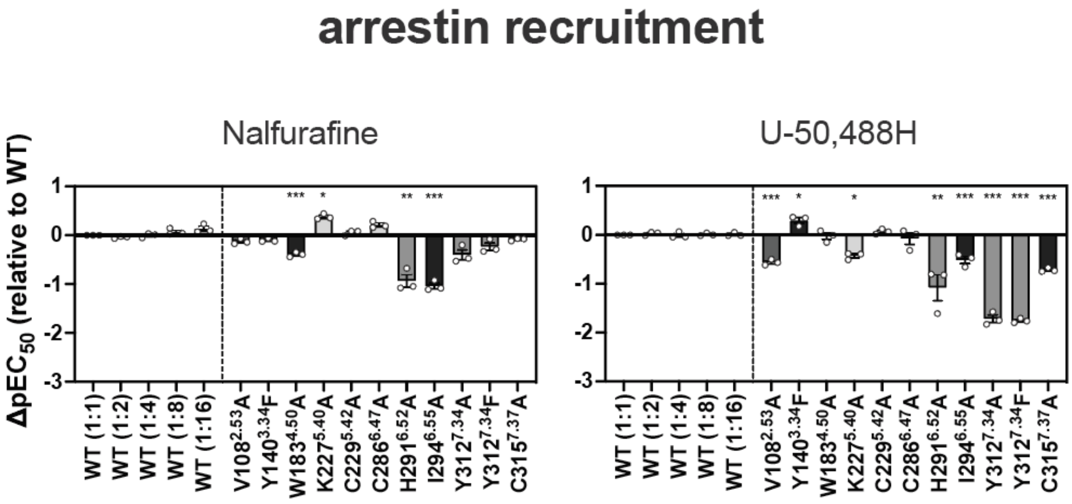


**Supplemental Figure 8. β-arrestin-recruiting activity analyzed by the NanoBiT-β-arrestin recruitment assay.**

Pharmacological parameters for the β-arrestin2-recruiting activity analyzed by the NanoBiT-β-arrestin recruitment assay. Data are presented as mean values ± SEM (*n* = 3-4; dots). For the individual experiments performed in parallel, data were normalized to the wild-type (WT) KOR (1:1) and presented as $\Delta$ $pEC_{50}$. The colors in the mutant bars correspond to the expression-matched WT conditions. Statistical analyses were performed using the ordinary one-way ANOVA followed by Dunnett tests with the expression-matched (colored) WT response. ns, $p > 0.05$; *$p < 0.05$; **$p < 0.01$; ***$p < 0.001$. Data are presented as mean values ± SEM (n=3; dots).

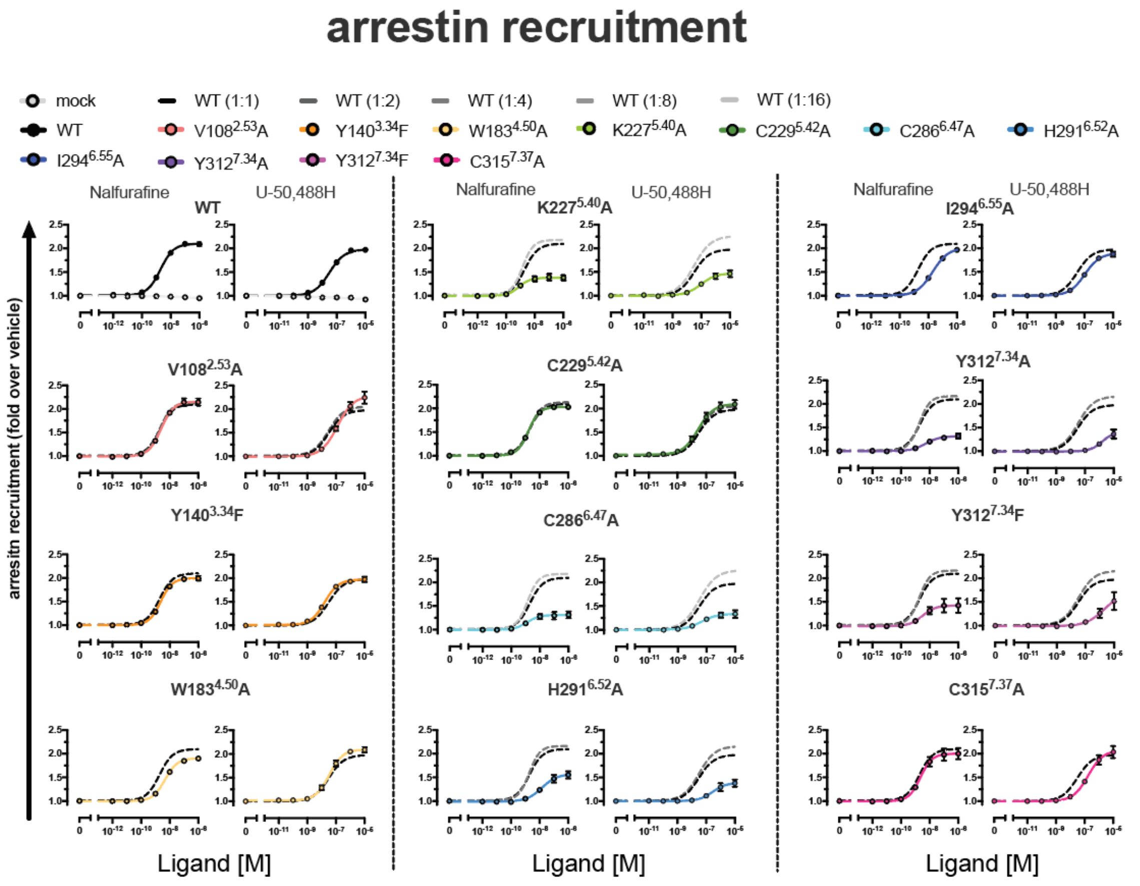


**Supplemental Figure 9. Concentration-response curves of the NanoBiT-β-arrestin recruitment assay.**

Dashed lines in the mutant panels represent the wild-type (WT) KOR (1:1, 1:2, 1:4, 1:8, or 1:16) response. Data are presented as mean values ± SEM (*n* = 3). Note that in numerous data points, the error bars are smaller than the size of the symbols, making them not visible.

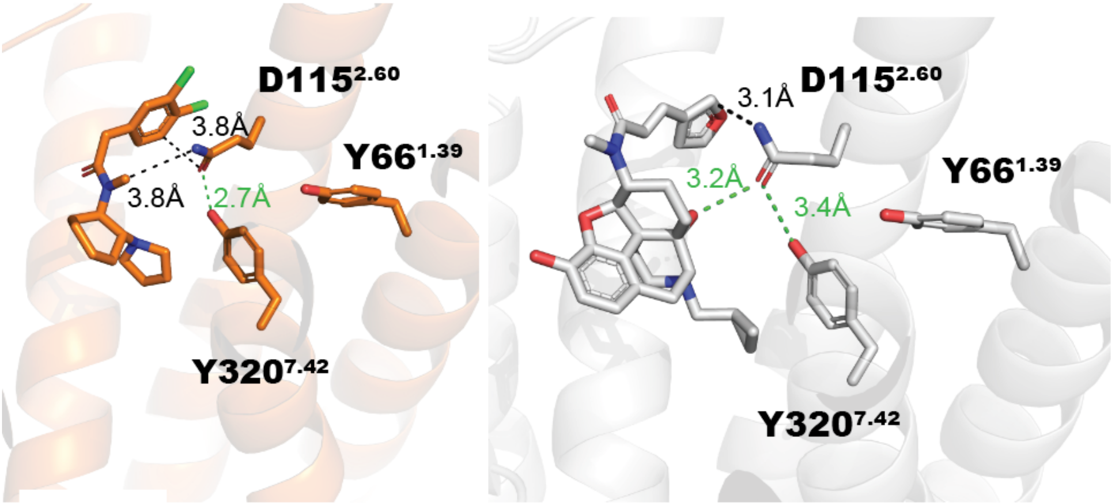


**Supplemental Figure 10. Different binding modes of D115$^{2.60}$ and ligand in U-50,488H (orange) - and nalfurafine(gray)-bound KOR-G$_i$ signaling complexes.**

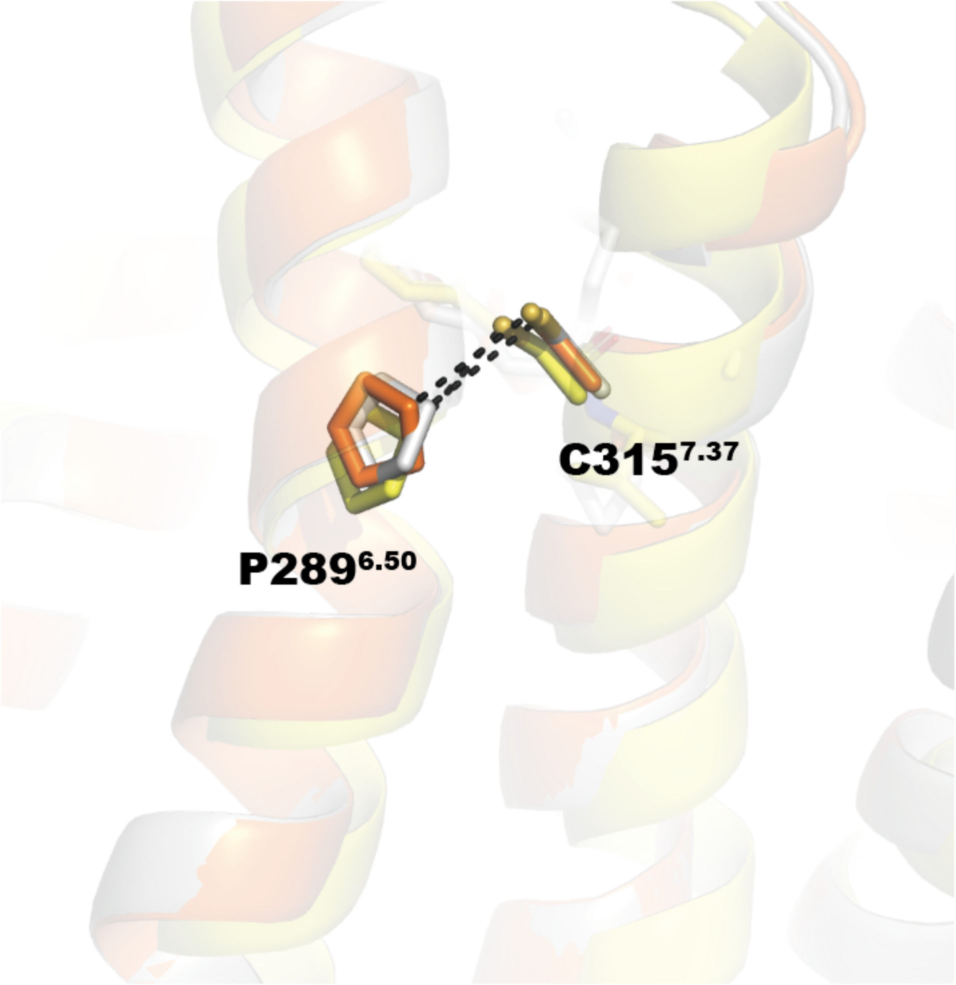


**Supplemental Figure 11. Interaction between the sidechains of P289$^{6.50}$ and C315$^{7.37}$.** Nalfurafine-bound KORs are in gray, U-50,488H-bound KORs in orange and inverse agonist JDTic-bound KORs in yellow.

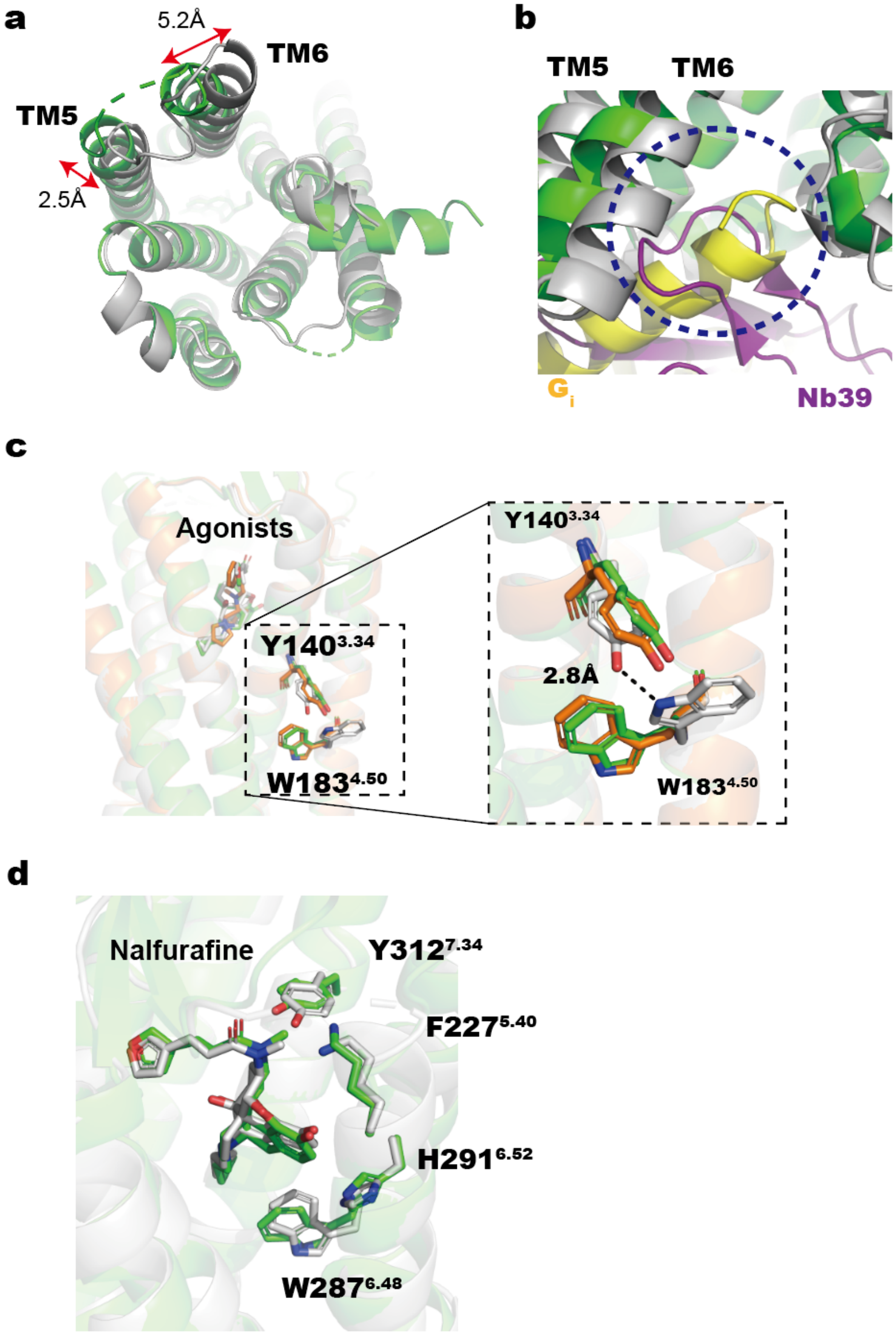
a
5.2Å
TM6
TM5
2.5Å
b
TM5
TM6
Gi
Nb39
c
Agonists
Y140 3.34
W183 4.50
Y140 3.34
2.8Å
W183 4.50
d
Nalfurafine
Y312 7.34
F227 5.40
H291 6.52
W287 6.48

**Supplemental Figure 12. Structural comparison of KOR-Nb39 complex and KOR-G protein complex in the nalfurafine-bound state.**

Superimposed view of the receptor region of the KOR-$G_i$ signaling complex (gray) and the KOR-Nb39 complex (green) in the nalfurafine-bound state from the intracellular side (**a**) and magnified view of the c-terminal helix of the G protein inserting into KOR and the binding site of Nb39 (**b**). Differences in the orientation of the side chains of Y140$^{3.34}$ and W183$^{4.50}$ are seen in the superposition diagram of the KOR-$G_i$ signaling complex bound with nalfurafine and U-50,488H (orange) and the KOR-Nb39 complex bound with nalfurafine (c). Different orientations of the side chains of amino acids involved in arrestin recruitment activity found in this study (d). In the nalfurafine-bound KOR-Gi signalling complex, the space between interacting amino acids is indicated by a black dotted line.

**Supplementary Table 1. Pharmacological parameters for the $G_i$-coupling activity and β-arrestin-recruiting activity.**

Pharmacological parameters for the $G_i$-coupling activity analyzed by the NanoBiT-G-protein dissociation assay and β-arrestin2-recruiting activity analyzed by the NanoBiT-β-arrestin recruitment assay. Data are presented as mean values ± SEM (*n* = 3-4; dots). For the individual experiments performed in parallel, data were normalized to the wild-type (WT) KOR (1:1) and presented as $E_{max}$ and $\Delta pEC_{50}$.

$G_i$ dissociation assay

| | Nalfurafine | | | | U-50,488H | | | |
|---|---|---|---|---|---|---|---|---|
| | Emax (%WT) | | pEC50 | | Emax (%WT) | | pEC50 | |
| | Mean | SEM | Mean | SEM | Mean | SEM | Mean | SEM |
| WT(1:1) | 100.0 | 0.0 | 10.22 | 0.21 | 100.0 | 0.0 | 8.69 | 0.08 |
| WT(1:2) | 100.0 | 1.0 | 10.11 | 0.11 | 100.4 | 0.7 | 8.63 | 0.05 |
| WT(1:4) | 98.3 | 0.8 | 9.96 | 0.12 | 97.6 | 1.6 | 8.42 | 0.02 |
| WT(1:8) | 90.5 | 2.1 | 9.86 | 0.08 | 93.2 | 0.8 | 8.14 | 0.02 |
| WT(1:16) | 77.8 | 1.0 | 9.83 | 0.07 | 82.4 | 0.1 | 7.92 | 0.06 |
| V108A | 102.8 | 1.1 | 9.95 | 0.08 | 97.6 | 1.4 | 8.48 | 0.09 |
| Y140F | 93.1 | 1.0 | 10.05 | 0.07 | 88.6 | 1.3 | 9.23 | 0.10 |
| W183A | 101.1 | 2.5 | 9.62 | 0.06 | 97.6 | 2.0 | 8.93 | 0.09 |
| K227A | 101.7 | 0.6 | 10.08 | 0.03 | 101.9 | 0.4 | 8.11 | 0.08 |
| C229A | 101.4 | 0.7 | 10.07 | 0.04 | 101.6 | 1.5 | 8.94 | 0.07 |
| C286A | 99.0 | 1.1 | 9.74 | 0.05 | 96.8 | 0.4 | 8.39 | 0.06 |
| H291A | 93.6 | 0.3 | 9.33 | 0.10 | 93.3 | 1.9 | 7.36 | 0.07 |
| I294A | 92.6 | 1.5 | 9.32 | 0.36 | 92.7 | 1.8 | 7.69 | 0.28 |
| Y312A | 95.8 | 3.8 | 8.90 | 0.09 | 100.1 | 2.4 | 6.73 | 0.22 |
| Y312F | 103.0 | 1.8 | 9.56 | 0.02 | 104.3 | 1.8 | 7.46 | 0.18 |
| C315A | 103.1 | 1.1 | 9.99 | 0.18 | 102.9 | 1.3 | 8.05 | 0.05 |

β-arrestin 2 recruitment assay

| | Nalfurafine | | | | U-50,488H | | | |
|---|---|---|---|---|---|---|---|---|
| | Emax (%WT) | | pEC50 | | Emax (%WT) | | pEC50 | |
| | Mean | SEM | Mean | SEM | Mean | SEM | Mean | SEM |
| WT(1:1) | 100.0 | 0.0 | 8.73 | 0.05 | 100.0 | 0.0 | 7.51 | 0.12 |
| WT(1:2) | 101.7 | 4.2 | 8.69 | 0.06 | 103.4 | 4.5 | 7.53 | 0.14 |
| WT(1:4) | 104.2 | 6.8 | 8.73 | 0.03 | 114.1 | 4.7 | 7.50 | 0.09 |
| WT(1:8) | 100.0 | 10.2 | 8.80 | 0.01 | 111.8 | 13.0 | 7.51 | 0.13 |
| WT(1:16) | 105.4 | 13.4 | 8.86 | 0.01 | 123.5 | 16.3 | 7.51 | 0.13 |
| V108A | 103.7 | 4.3 | 8.60 | 0.02 | 124.4 | 6.7 | 6.94 | 0.10 |
| Y140F | 89.2 | 3.2 | 8.63 | 0.06 | 95.0 | 6.0 | 7.80 | 0.08 |
| W183A | 80.9 | 3.1 | 8.34 | 0.07 | 107.0 | 3.2 | 7.48 | 0.17 |
| K227A | 34.1 | 3.0 | 9.11 | 0.01 | 46.0 | 5.5 | 7.08 | 0.12 |
| C229A | 93.2 | 3.4 | 8.78 | 0.04 | 105.3 | 4.4 | 7.57 | 0.14 |
| C286A | 26.9 | 5.3 | 8.94 | 0.08 | 32.4 | 6.7 | 7.43 | 0.15 |
| H291A | 51.8 | 4.1 | 7.80 | 0.08 | 39.2 | 4.1 | 6.43 | 0.18 |
| I294A | 91.5 | 4.9 | 7.69 | 0.01 | 87.3 | 1.8 | 6.99 | 0.06 |
| Y312A | 28.3 | 3.2 | 8.33 | 0.13 | 41.3 | 9.8 | 5.79 | 0.05 |
| Y312F | 38.7 | 11.3 | 8.50 | 0.10 | 66.1 | 21.2 | 5.75 | 0.10 |
| C315A | 89.7 | 5.5 | 8.64 | 0.04 | 104.3 | 8.0 | 6.79 | 0.10 |

**Supplementary Table 2 Cryo-EM data collection, refinement, and validation statistics**

| | U-50,488H bound KOR-$G_i$ signaling complex (EMD-38751) (PDB: 8XXJ) | Nalfurafine bound KOR-$G_i$ signaling complex (EMD-37545) (PDB: 8WHR) |
|---|---|---|
| **Data collection and processing** | | |
| Magnification | 105,000 | 105,000 |
| Voltage (keV) | 300 | 300 |
| Electron exposure ($e^-/Å^2$) | 60 | 60 |
| Defocus range (μm) | -0.7 to -1.5 | -0.7 to -1.5 |
| Pixel size (Å) | 0.675 | 0.675 |
| Symmetry imposed | C1 | C1 |
| Initial particle images (no.) | 4,148,764 | 7,066,947 |
| Final particle images (no.) | 1,225,096 | 858,423 |
| Map resolution (Å) | 2.90 | 2.76 |
| FSC threshold | 0.143 | 0.143 |
| Map resolution range (Å) | 2.47-14.99 | 2.36-6.67 |
| **Refinement** | | |
| Initial model used (PDB code) | 7YIT(KOR), 6CMO ($G\alpha_i$, scFv16), 6DDE (Gβ, Gγ) | 7YIT(KOR), 6CMO ($G\alpha_i$, scFv16), 6DDE (Gβ, Gγ) |
| Model resolution (Å) | 2.90 | 2.76 |
| FSC threshold | 0.5 | 0.5 |
| Map sharpening *B* factor ($Å^2$) | 128.4 | 110.4 |
| Model composition | | |
| Non-hydrogen atoms | 8,284 | 8,358 |
| Protein residues | 1,103 | 1,107 |
| Ligands | 1 | 1 |
| *B* factors ($Å^2$) | | |
| Protein | 71.10 | 58.51 |
| Ligand | 112.45 | 80.37 |
| R.m.s. deviations | | |
| Bond lengths (Å) | 0.003 | 0.003 |
| Bond angles (°) | 0.531 | 0.492 |

| | | |
|---|---|---|
| Validation | | |
| MolProbity score | 2.37 | 1.93 |
| Clashscore | 9.63 | 6.11 |
| Poor rotamers (%) | 8.47 | 4.05 |
| Ramachandran plot | | |
| Favored (%) | 97.13 | 97.33 |
| Allowed (%) | 2.87 | 2.67 |
| Disallowed (%) | 0 | 0 |
